# tRNA modification enzyme-dependent redox homeostasis regulates synapse formation and memory

**DOI:** 10.1101/2023.11.14.566895

**Authors:** Kimberly R. Madhwani, Shanzeh Sayied, Carlson H. Ogata, Caley A. Hogan, Jenna M. Lentini, Moushami Mallik, Jennifer L. Dumouchel, Erik Storkebaum, Dragony Fu, Kate M. O’Connor-Giles

**Affiliations:** Neuroscience Graduate Program, Brown University, Providence, RI, USA; Department of Neuroscience, Brown University, Providence, RI, USA; Department of Biology, Brown University, Providence, RI, USA; Laboratory of Genetics, University of Wisconsin-Madison, Madison, WI, USA; Department of Biology, Center for RNA Biology, University of Rochester, Rochester, NY, USA; Molecular Neurobiology Laboratory, Donders Institute for Brain, Cognition, and Behaviour, Radboud University, Nijmegen, NL; Therapeutic Sciences Graduate Program, Brown University, Providence, RI, USA; Carney Institute for Brain Sciences, Brown University, Providence, RI, USA

**Keywords:** RNA modification, tRNA methyltransferase, mRNA translation, oxidative stress, neurodevelopment

## Abstract

Post-transcriptional modification of RNA regulates gene expression at multiple levels. ALKBH8 is a tRNA modifying enzyme that methylates wobble uridines in specific tRNAs to modulate translation. Through methylation of tRNA-selenocysteine, ALKBH8 promotes selenoprotein synthesis and regulates redox homeostasis. Pathogenic variants in ALKBH8 have been linked to intellectual disability disorders in the human population, but the role of ALKBH8 in the nervous system is unknown. Through *in vivo* studies in *Drosophila*, we show that ALKBH8 controls oxidative stress in the brain to restrain synaptic growth and support learning and memory. *ALKBH8* null animals lack wobble uridine methylation and exhibit a global reduction in protein synthesis, including a specific decrease in selenoprotein levels. Loss of *ALKBH8* or independent disruption of selenoprotein synthesis results in ectopic synapse formation. Genetic expression of antioxidant enzymes fully suppresses synaptic overgrowth in *ALKBH8* null animals, confirming oxidative stress as the underlying cause of dysregulation. *ALKBH8* animals also exhibit associative learning and memory impairments that are reversed by pharmacological antioxidant treatment. Together, these findings demonstrate the critical role of tRNA modification in redox homeostasis in the nervous system and reveal antioxidants as a potential therapy for ALKBH8-associated intellectual disability.

**Significance Statement:** tRNA modifying enzymes are emerging as important regulators of nervous system development and function due to their growing links to neurological disorders. Yet, their roles in the nervous system remain largely elusive. Through *in vivo* studies in *Drosophila*, we link tRNA methyltransferase-regulated selenoprotein synthesis to synapse development and associative memory. These findings demonstrate the key role of tRNA modifiers in redox homeostasis during nervous system development and highlight the potential therapeutic benefit of antioxidant-based therapies for cognitive disorders linked to dysregulation of tRNA modification.

## Introduction

Nervous system development relies on the dynamic regulation of gene expression at the levels of transcription, translation, and protein stability (1–9). Neuronal cell fate diversification, memory consolidation, and neuronal survival all depend on the tight regulation of translation to coordinate protein expression and control protein levels (10–12). tRNAs are central components of the translational machinery that decode mRNA by linking codons and the corresponding amino acids (13, 14). Post-transcriptional modification of RNA is emerging as a critical regulator of protein synthesis in the nervous system (15–18). tRNAs are the most heavily modified cellular RNA, with more than 100 known modifications, modulating protein folding, stability, and function (19–26). Increasing links between mutations in tRNAs and neurological disease highlight the importance of understanding tRNA processing and function in the context of the nervous system (15–18, 27–31). However, it remains unclear why disruption of tRNA biogenesis and function disproportionately affects the nervous system, underscoring the need for further investigation.

Uridines at the wobble position of tRNA anticodon loops are almost always modified in all species. The methylation and thiolation of tRNA wobble uridines can modulate decoding of cognate codons versus codons with non-standard Watson-Crick base pairing (32, 33). In yeast, a subset of wobble uridines is methylated by tRNA methyltransferase 9 (Trm9) to generate the 5-methoxycarbonyl-methyluridine (mcm^5^U) modification (34). Trm9-dependent tRNA modifications promote the expression of mRNAs enriched for specific codons that encode DNA damage response proteins (35–38). Consistently, Trm9-deficient yeast cells exhibit increased sensitivity to DNA damaging agents. Thus, disruption of wobble uridine methylation can have far-reaching effects on protein expression and cellular physiology.

Animals and plants express orthologs of yeast Trm9 that are encoded by the AlkB homolog 8 (ALKBH8) and tRNA methyltransferase 9B (TRMT9B) genes (39–45). ALKBH8 and TRMT9B contain a Class I *S*-adenosyl-L-methionine (SAM)-dependent methyltransferase domain, similar to yeast Trm9. In addition, ALKBH8 contains an RNA recognition binding motif and an active 2- oxoglutarate-Fe(II) oxygenase domain (40, 45). While the substrates of TRMT9B modification remain uncertain, studies in mammals and flies have demonstrated that ALKBH8 carries out the canonical yeast Trm9 function of methylating tRNA wobble uridines (39, 41, 44). In mammalian cells, ALKBH8 methylates wobble uridines in tRNA-Selenocysteine-UGA, tRNA-Glutamate-UUC, tRNA-Arginine-UCU, and tRNA-Glycine-UCC (39, 44, 46). Moreover, the 2-oxoglutarate-Fe(II) oxygenase domain of ALKBH8 has been shown to catalyze the hydroxyl modification of tRNA-Glycine-UCC (40, 43, 45).

Loss of ALKBH8 and its associated methylation disrupts DNA repair and oxidative stress responses as well as cellular senescence and mitochondrial activity in mammalian cells (47–52). ALKBH8 regulates oxidative stress via regulation of selenoprotein synthesis, which relies on methylation of tRNA-selenocysteine (tRNA^Sec^) for recoding UGA stop codons. Selenoproteins are potent antioxidants and play a key role in controlling cellular ROS levels. Notably, mutations in selenoprotein biosynthesis machinery are associated with neurodevelopmental disorders (53–57). More recently, mutations in *ALKBH8* have been identified as the underlying cause of autosomal-recessive intellectual disability in multiple families (58–61). These findings reveal a critical requirement for ALKBH8 in nervous system development and function. However, ALKBH8 function in the nervous systems remains unexplored.

To investigate the role of ALKBH8 in nervous system development and function *in vivo*, we generated null alleles in *Drosophila*. Loss of *ALKBH8*-dependent tRNA methylation results in reduced selenoprotein synthesis and ectopic synapse formation. We find that synaptic overgrowth is suppressed by genetic expression of antioxidant enzymes, implicating oxidative stress in the dysregulation of synaptogenesis in the absence of ALKBH8. We also find that loss of *ALKBH8* impairs learning and memory, consistent with the cognitive impairments observed in patients with ALKBH8-associated intellectual disability. Remarkably, pharmacological antioxidant treatment fully suppresses learning and memory deficits. Together, our results uncover a neuronal role for ALKBH8 in redox homeostasis, synapse formation, and circuit function, providing insight into the cellular mechanisms underlying ALKBH8-associated intellectual disability and potential treatments.

## Results

### ALKBH8 attenuates synapse formation

To determine the neurodevelopmental role of ALKBH8 *in vivo*, we generated null alleles of *Drosophila ALKBH8* (CG17807) using CRISPR-based gene editing (62, 63). Briefly, an interfering cassette encoding a visible marker flanked by piggyBac inverted repeat sequences was inserted immediately downstream of the *ALKBH8* translation start site, generating a null allele, *ALKBH8*^1-IC^ (IC, interfering cassette; Figure 1A) (41). To control for potential off-target cleavage, we created an independent null allele, *ALKBH8*^2-IC^, using a distinct gRNA (Figure 1A). To measure synapse formation in the absence of ALKBH8, we turned to the well-characterized, highly stereotyped glutamatergic *Drosophila* larval neuromuscular junction 4 (NMJ4) and quantified synaptic boutons, the axonal varicosities where synapses form (64). Loss of *ALKBH8* resulted in synaptic overgrowth, with a 51% increase in bouton number, indicating that ALKBH8 restrains synapse formation (Figure 1B,C,E). Restoration of ALKBH8 expression under endogenous regulation by removal of the interfering cassette restored synapse formation to near control levels (Figure 1D,E; *ALKBH8*^1-ER^, ER, endogenous rescue). We observed similar overgrowth and full genetic rescue in *ALKBH8*^2-IC^ and *ALKBH8*^2-ER^, respectively (Figure 1F-I). These results indicate that ALKBH8 acts as a brake on synapse formation.

**Figure 1.**
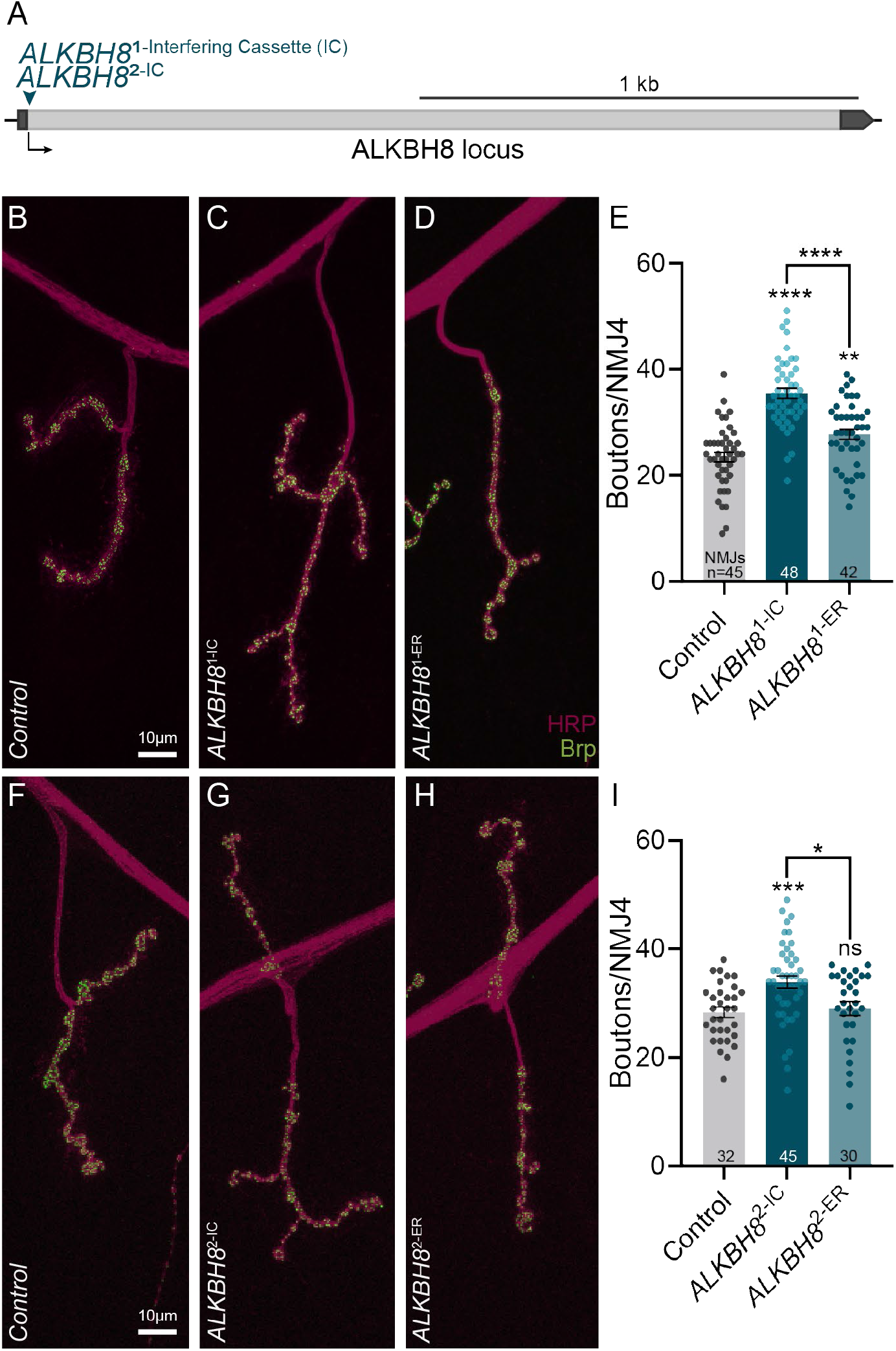
ALKBH8 attenuates synapse formation. (**A**) Schematic illustrating the *ALKBH8* locus with alleles generated in this study. The teal triangle marks the site of interfering cassette insertion. (**B-I**) Representative confocal images of NMJ4 in indicated genotypes labeled with the neuronal membrane marker HRP (magenta) and active zone marker Brp (green; **B-D, F-H**) and NMJ4 bouton quantification (**E,I**). Scale bar: 10 µm. Data points represent individual NMJs from 30-48 animals with n indicated in graph. One-way ANOVA followed by Dunnett’s T3 multiple comparisons test. Data is represented as mean ± SEM. \*\*\*\**p* < 0.0001, \*\*\**p* = 0.0007, \*\**p* = 0.005, \**p* = 0.015, *ns* = not significant.

### Loss of ALKBH8 abolishes wobble uridine methylation and hydroxylation

Post-transcriptional modification of wobble uridines at position 34 of the tRNA anticodon loop (U_34_) is observed in all tested species. U_34_ modifications rely on multiple enzymatic pathways that have primarily been studied in yeast, where they modulate translation based on codon usage (17, 33, 37). Here, we focus on one branch of the U_34_ modification pathway in which the multi-protein Elongator complex first methylates U_34_ to generate 5-carbonylmethyluridine (cm^5^U; Figure 2A, orange). In mammals, ALKBH8 then adds a second methyl group to yield 5- methoxycarbonyl-methyluridine (mcm^5^U; Figure 2A, teal). Further modification by the Ncs2-Ncs6 thiolase complex generates a terminal 5-methoxycarbonylmethyl-2-thiouridine modification (mcm^5^s^2^U, Figure 2A, magenta). ALKBH8 also hydroxylates mcm^5^U in tRNA-Glycine-UCC to form 5-methoxycarbonylhydroxymethyluridine (mchm^5^U, Figure 2A, green) (40, 44).

**Figure 2.**
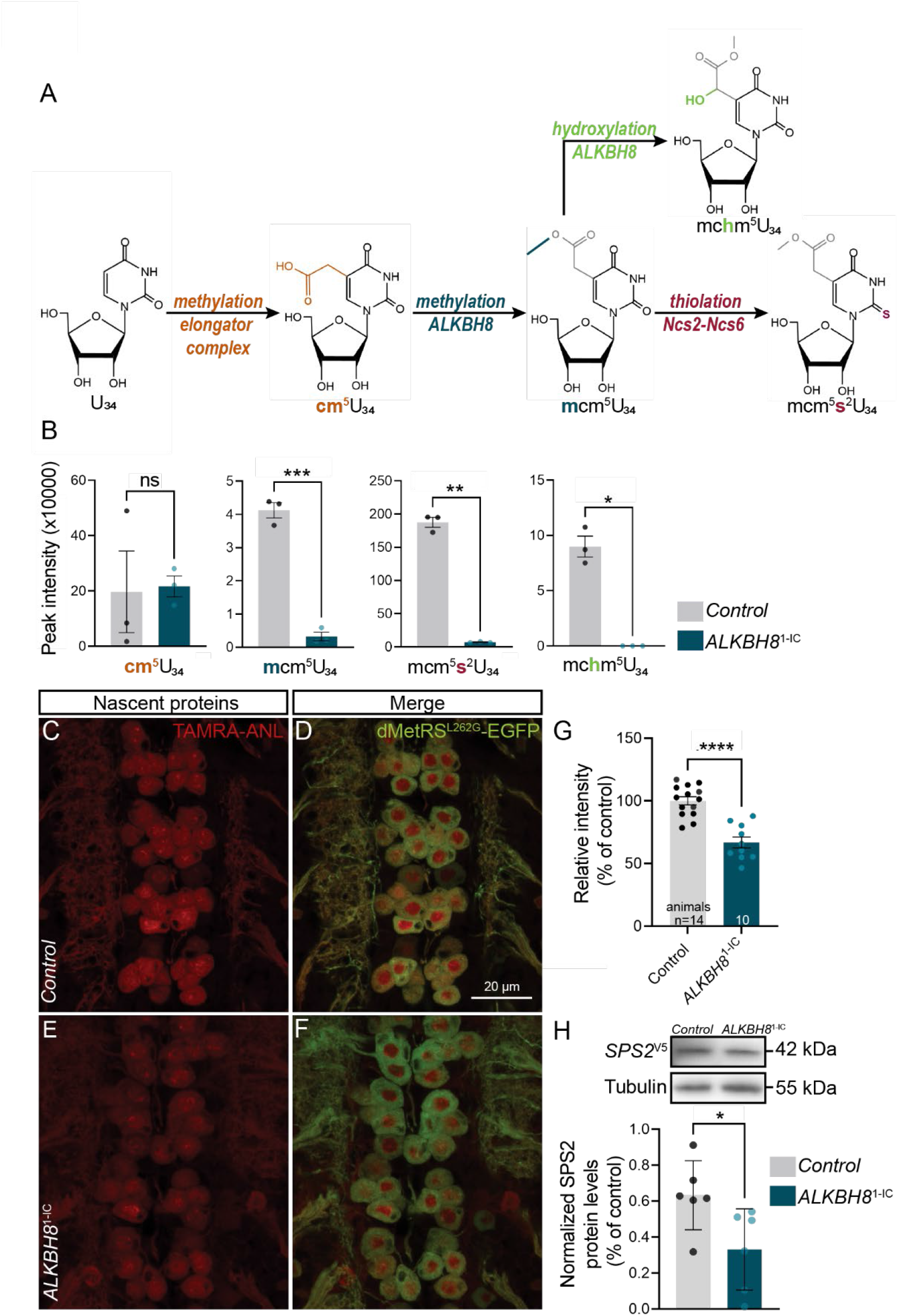
ALKBH8 methylates tRNA wobble uridines in the brain and promotes global translation and selenoprotein synthesis. (**A**) Schematic illustrating the ALKBH8 enzymatic pathway. (**B**) Peak area of modified wobble uridines normalized to the canonical nucleosides A, U, G, and C in nucleosides isolated from control and *ALKBH8*^1-IC^ adult heads. 3 biological replicates per genotype. (**C-F**) Representative images of FUNCAT labelling in larval motor neurons of control and *ALKBH8*^1-IC^ animals expressing *dMetRS^L262G^-EGFP*. Scale bar: 20 µm. (**G**) Quantification of FUNCAT (TAMRA-ANL) signal intensity. Each point denotes the average signal intensity of a single motor neuron cluster per animal with n indicated in graph. (**H**) Endogenous tagging of SPS2 with the small peptide tag V5 (*SPS2*^V5^) shows a reduction in SPS2 protein levels in the absence of *ALKBH8*. Representative blot probing SPS2 protein levels in control and *ALKBH8*^1-IC^ larval ventral nerve cord/body walls. Tubulin was used as a loading control and SPS2 levels were normalized to tubulin. 6 biological replicates per genotype. Welch’s t-test (B,H), Unpaired t-test (G). Data is represented as mean ± SEM. \*\*\*\**p* < 0.0001, \*\*\**p* = 0.0005, \*\**p* = 0.0016, \**p* < 0.05, *ns* = not significant.

To evaluate the enzymatic role of ALKBH8 in *Drosophila*, we monitored wobble uridine modification status in control and *ALKBH8*^1-IC^ animals using liquid chromatography-mass spectrometry (LC-MS/MS) of digested RNA from the brain. Total RNA isolated from heads of mated control and *ALKBH8*^1-IC^ flies was nuclease digested and dephosphorylated to produce individual nucleosides followed by LC-MS analysis of modifications (39, 65). We observed a near-complete loss of wobble uridine methylation and the dependent downstream thiolation and hydroxylation modifications in *ALKBH8*^1-IC^ animals (Figure 2B). No significant changes in other modifications were detected in the *ALKBH8* null animals (Figure S1). These results demonstrate that ALKBH8 is required for tRNA wobble uridine modification in *Drosophila*.

### ALKBH8 promotes global and selenoprotein translation

Post-transcriptional modification of tRNAs plays a critical role in protein synthesis. Thus, we investigated a potential function for ALKBH8 in protein translation in the nervous system. To directly assess global protein synthesis *in vivo*, we used fluorescent noncanonical amino-acid tagging (FUNCAT) (66, 67). This technique employs a mutated methionyl-tRNA synthetase tagged with green fluorescent protein (GFP) that can be expressed in a cell-specific manner using the UAS-Gal4 system (*UAS-MetRS*^L262G^*-EGFP*). The mutated methionyl-tRNA synthetase enables the incorporation of the non-canonical amino acid azidonorleucine (ANL) into nascent proteins, allowing the subsequent tagging of newly synthesized proteins with a red-fluorescent dye tetramethylrhodamine (TAMRA) by click chemistry, followed by visualization using confocal microscopy. In this system, the rate of protein synthesis is proportional to fluorescent labelling intensity. We used *OK371-Gal4* to drive *dMetRS^L262G^-EGFP* in control and *ALKBH8*^1-IC^ larval motor neurons and evaluated protein synthesis rates. Newly synthesized protein levels were ∼33% lower in *ALKBH8*^1-IC^ motor neurons than control as measured by TAMRA-ANL intensity (Figure 2C-G). This finding demonstrates that loss of *ALKBH8* inhibits global protein translation in motor neurons *in vivo*.

ALKBH8 plays a key role in the synthesis of selenocysteine-containing selenoproteins through the methylation of tRNA^Sec^ (44, 47, 49, 51). tRNA^Sec^ is the only tRNA with a terminal mcm^5^U modification. Methylation of tRNA^Sec^ catalyzes the incorporation of the rare 21^st^ amino acid selenocysteine, an amino acid not formally designated in the genetic code but instead through the recoding of in-frame UGA stop codons (68). Humans express 25 selenoproteins, of which three are conserved in *Drosophila* (68–71). Of the three *Drosophila* selenoproteins, selenophosphate synthetase 2 (SPS2) is broadly expressed and, intriguingly, is not only a selenoprotein but also performs a key role in selenocysteine biosynthesis by promoting the formation of the selenium donor, selenophosphate (72, 73). Thus, loss of SPS2 impairs all selenoprotein synthesis.

To determine the role of ALKBH8 in selenoprotein synthesis in *Drosophila*, we engineered a V5 peptide-tagged *SPS2* allele using CRISPR (*SPS2*^V5^). We then measured SPS2 levels in control and *ALKBH8*^1-IC^ larval central nervous system/body wall preparations. We observed a 48% decrease in SPS2 protein levels in the absence of ALKBH8, demonstrating a conserved role for ALKBH8 in promoting selenoprotein expression (Figure 2H).

### Selenoproteins restrain synaptic growth

Selenoproteins are key regulators of oxidative stress (68, 71, 74). Intriguingly, excess oxidative stress is known to cause synaptic overgrowth. Finding that ALKBH8 modifies tRNA^Sec^, promotes selenoprotein expression in the brain and attenuates synapse formation, we next asked if selenoproteins regulate synaptic growth. SPS2 is essential for selenoprotein biosynthesis, thus we generated a null allele*, SPS2*^1^, and obtained an independent insertion allele, *SPS2*^EY00228^ (75). We quantified synaptic bouton number in three mutant combinations and observed synaptic overgrowth similar to *ALKBH8*, revealing a role for selenoproteins in synaptogenesis (Figure 3A-E,G).

**Figure 3.**
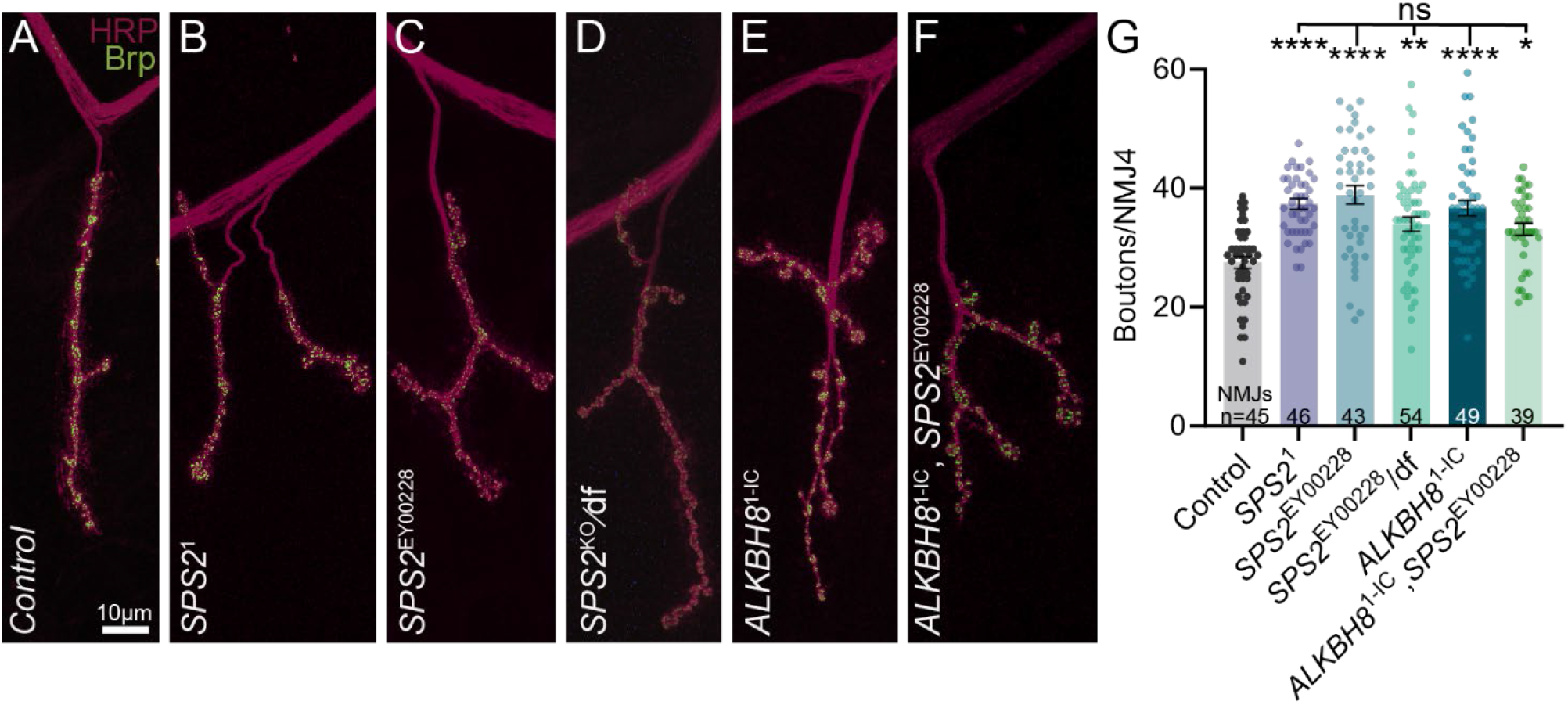
Selenoproteins regulate synapse formation. (**A-F**) Representative confocal images of NMJ4 in indicated genotypes labeled with the neuronal membrane marker HRP (magenta) and active zone marker Brp (green). Scale bar: 10 µm. (**G**) NMJ4 bouton quantification. Data points represent individual NMJs from 39-54 animals with n indicated in graph. Kruskal-Wallis test followed by Dunn’s multiple comparisons test. Data is represented as mean ± SEM. \*\*\*\**p* < 0.0001, \*\**p* = 0.0018, \**p* = 0.021, *ns* = not significant.

A role for selenoproteins in synapse formation suggests a model in which ALKBH8 regulates synaptic growth through the regulation of selenoprotein synthesis. If ALKBH8 and SPS2 act together in a pathway to attenuate synaptic growth, loss of both *ALKBH8* and *SPS2* will not result in greater ectopic synapse formation than loss of either alone. Indeed, double mutants exhibit synaptic growth comparable to single *ALKBH8* and *SPS2* null animals, indicating that *ALKBH8* and *SPS2* act in the same pathway to regulate synapse development (Figure 3F,G).

### Antioxidant enzymes suppress ALKBH8-associated synaptic overgrowth

Our findings indicate a causal link between ALKBH8-dependent regulation of selenoprotein synthesis and synaptic overgrowth and suggest oxidative stress underlies synaptic overgrowth induced by loss of *ALKBH8*. We tested this model by assessing the ability of genetically encoded antioxidants to suppress ALKBH8-associated synaptic overgrowth.

Specifically, we ubiquitously overexpressed *catalase*, *superoxide dismutase 1* (SOD1), or *superoxide dismutase 2* (SOD2), which encode conserved ROS scavenging enzymes that act as a first line of defense against oxidative stress, in control and *ALKBH8*^1-IC^ animals using the UAS-Gal4 system. Catalase facilitates the breakdown of hydrogen peroxide (H_2_O_2_) into water and oxygen, while SOD1 and SOD2 catalyze the decomposition of superoxide radicals into H_2_O_2_ and water (76, 77). A previous study found that RNAi-mediated knockdown of SODs or catalase resulted in a significant increase in synaptic growth, demonstrating the important roles of antioxidant enzymes in synapse development (78). We find that overexpression of *catalase*, *SOD1*, or *SOD2* in a control background had no effect on synapse formation (Figure S2). In contrast, all three enzymes fully suppressed synaptic overgrowth in *ALKBH8*^1*-*IC^ animals (Figure 4A-F), consistent with the conclusion that oxidative stress is the cause of synaptic overgrowth.

**Figure 4.**
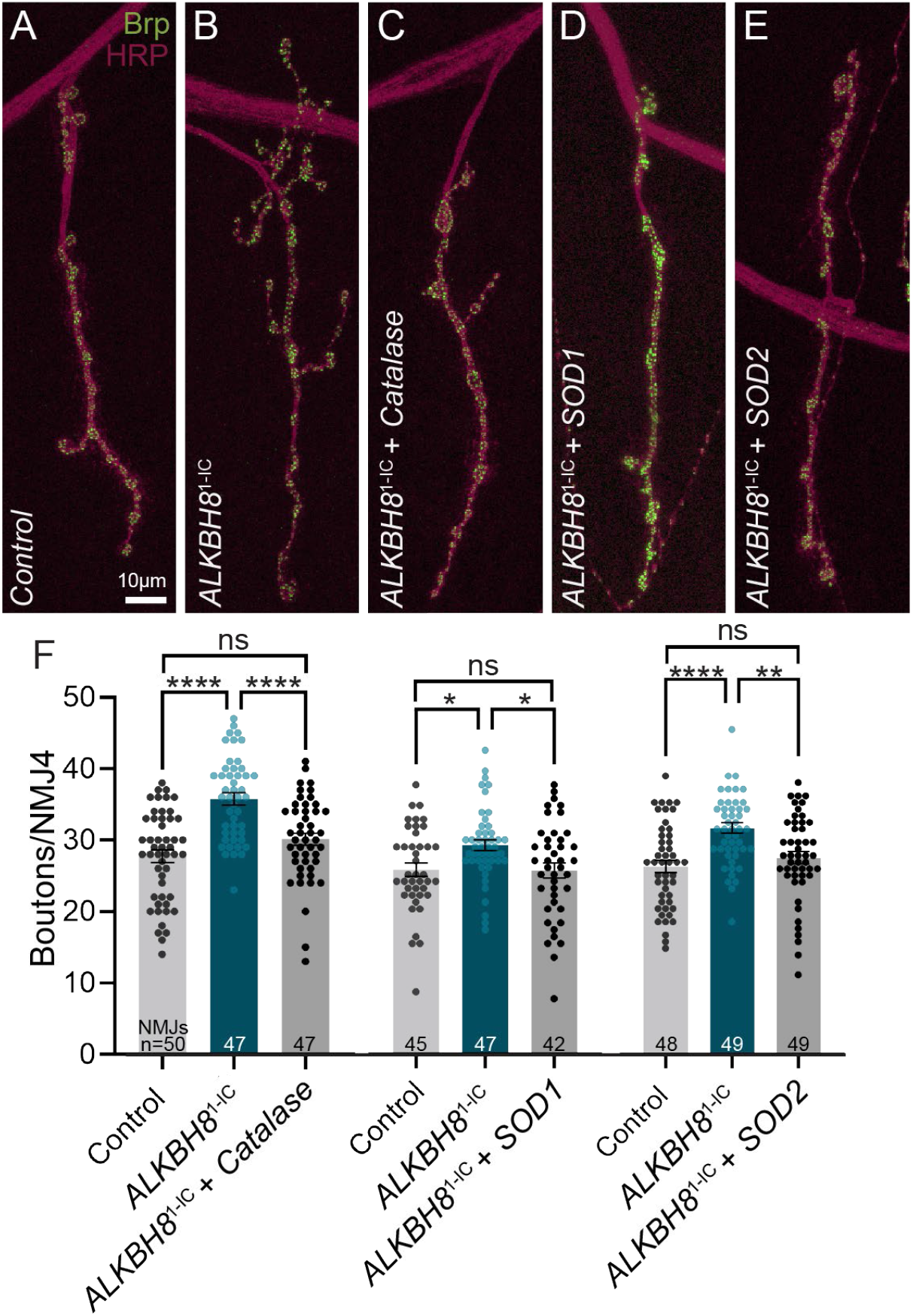
Antioxidant enzymes suppress synaptic overgrowth in *ALKBH8* mutants. (**A-E**) Representative confocal images of NMJ4 in indicated genotypes labeled with the neuronal membrane marker HRP (magenta) and active zone marker Brp (green). Scale bar: 10 µm. (**F**) Ubiquitous expression of antioxidant enzymes in *ALKBH8* null animals suppresses synaptic overgrowth. Data points represent individual NMJs from 42-50 animals with n indicated in graph. One-Way ANOVA followed by Dunnett’s T3 multiple comparisons test. Data is represented as mean ± SEM. \*\*\*\**p* < 0.0001, \*\**p* = 0.014, \**p* < 0.05, *ns* = not significant.

### ALKBH8 reduces oxidative stress

Selenoproteins are recruited when oxidative stress is greater than the cell’s basal capacity to maintain balance (79–81). To assess whether ALKBH8 provides protection against oxidative stressors, we exposed control and *ALKBH8*^1-IC^ adult flies to hydrogen peroxide (H_2_O_2_) or paraquat. H_2_O_2_ functions as a messenger in redox signaling and its dysregulation can impede oxidative stress response signaling cascades (82–85). Acting through a distinct mechanism, paraquat is a herbicide and neurotoxin that increases free radicals and impedes the function of mitochondrial complex I-III and complex II-III (86–89). We assessed mortality after 24-hours of exposure and found that animals lacking *ALKBH8* exhibit 23% or 53% greater mortality than control animals following H_2_O_2_ or paraquat exposure, respectively (Figure 5A). These findings demonstrate that ALKBH8 promotes systemic responses to oxidative stress.

**Figure 5.**
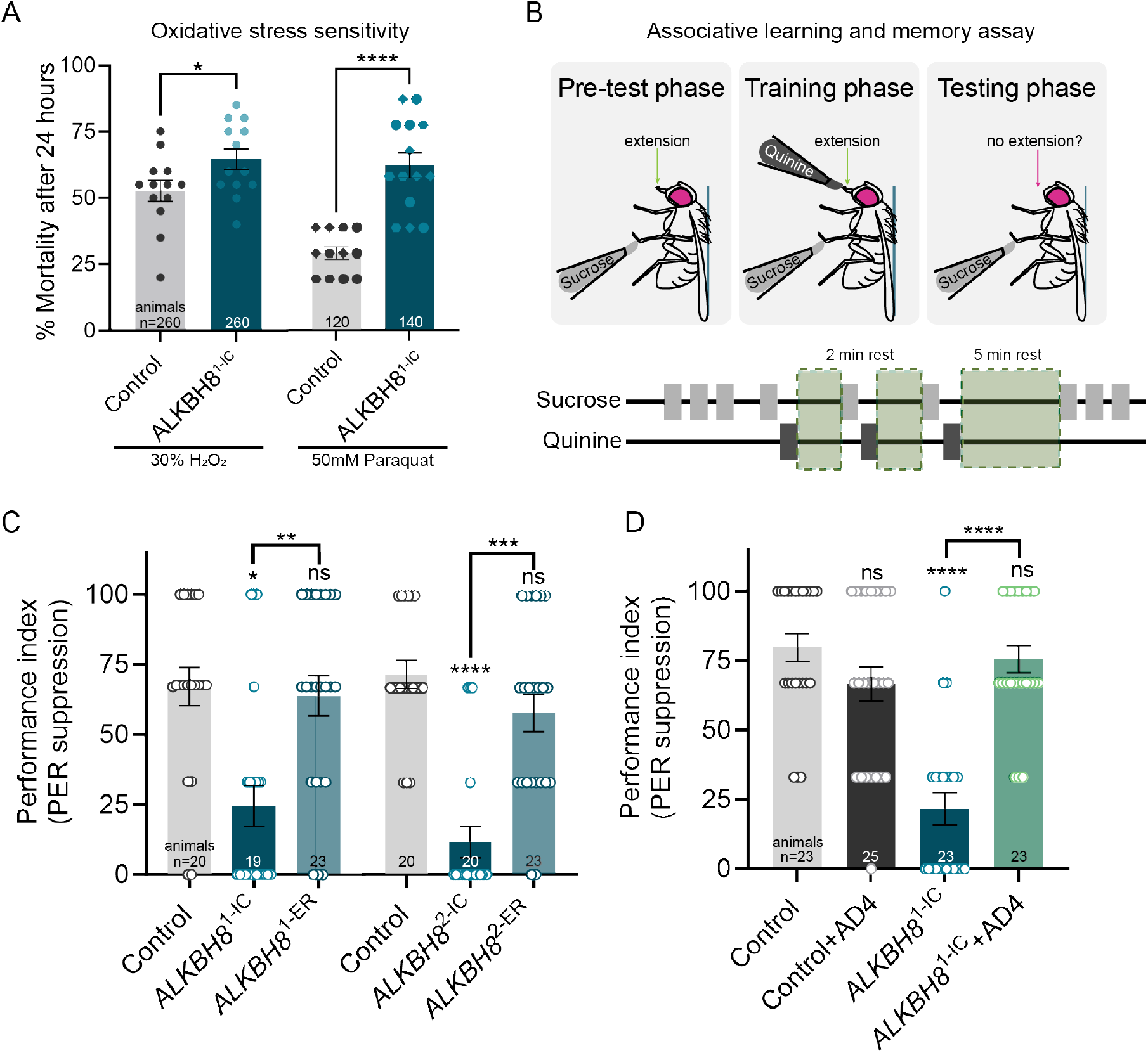
ALKBH8 promotes learning and memory by reducing oxidative stress. (**A**) Oxidative stress resistance is diminished in *ALKBH8*^1-IC^ animals. Each point denotes 20 or 10 3-5-day-old adult flies exposed to 30% H_2_O_2_ or 50mM paraquat, respectively, with n indicated in graph. (**B**) Schematic illustrating taste aversion assay adapted from Masek et al. 2015. (**C**) Quantification of Proboscis Extension Reflex (PER) suppression. Data points represent individual animals with n indicated in graph. (**D**) Raising *ALKBH8*^1-IC^ animals on food containing 40µg/ml AD4 (N-acetylcysteine amide) restores PER suppression to control levels. Data points represent individual animals with n indicated in graphs. Welch’s t-test (A), Kruskal-Wallis test followed by Dunn’s multiple comparisons test (C,D). Data is represented as mean ± SEM. \*\*\*\**p* < 0.0001, \*\*\**p* = 0.0002, \*\**p* < 0.01, \**p* < 0.05, *ns* = not significant.

### ALKBH8 promotes learning and memory by reducing oxidative stress

To date, two missense and three nonsense mutations, all in the last coding exon of *ALKBH8*, have been identified as the cause of autosomal-recessive intellectual disability (58–61). Three of these mutations result in a complete absence of the tRNA wobble uridine methylation modification in affected individuals. Due to these growing links between ALKBH8 and neurological disease, we explored the downstream behavioral consequences of the loss of *ALKBH8* using our animal model. We exploited the *Drosophila* gustatory system and a well-established experimental model to measure associative learning and memory (90, 91). Briefly, presentation of a sweet tastant to taste receptors on the tarsi (legs) of starved flies induces the Proboscis Extension Reflex (PER), a robust feeding behavior. To the formation of aversive taste memory, the sweet tastant sucrose is applied to the tarsi of starved flies while the bitter tastant quinine is simultaneously presented to the proboscis (mouth). Repeated training results in reduced PER response when a fly is subsequently presented with sucrose alone, indicating associative memory (Figure 5B) (92–95).

To ensure that the PER responses were not influenced by abnormal baseline taste preferences, we first measured PER by solely presenting sucrose or quinine to the tarsi. The performance index indicates how much PER was observed. As expected, control, *ALKBH8*^1-IC^ null, and *ALKBH8*^1-ER^ lines all showed a high PER response to sucrose and significantly lower PER response to quinine (Figure S3). When we assessed associative memory in *ALKBH8*^1-IC^ animals, we found that control animals suppressed proboscis extension following training, whereas *ALKBH8*^1-IC^ animals did not, indicating a severe deficit in associative memory. Memory was completely restored upon genetic re-expression of *ALKBH8*, revealing that ALKBH8 is required for learning and memory in this assay (Figure 5C). We corroborated this result by measuring learning and memory in our independent *ALKBH8* line, *ALKBH8*^2-IC^. Similarly, *ALKBH8*^2-IC^ animals exhibit an inability to retain memories while genetic restoration of *ALKBH8* under its endogenous promoter fully rescues this deficit (Figure 5C).

This finding prompted us to investigate whether antioxidant treatment can rescue learning and memory deficits in *ALKBH8* animals. To test this, we reared control and *ALKBH8*^1-IC^ animals on food supplemented with the antioxidant N-acetylcysteine amide (40µg/ml AD4 in water) or vehicle control (water) and evaluated learning performance in 7-day old adults. AD4-fed control animals displayed no significant learning differences compared to untreated control animals, whereas AD4 fully restored memory in *ALKBH8*^1-IC^ animals to control levels (Figure 5D). Collectively, these findings indicate that ALKBH8-dependent redox homeostasis regulates synapse formation and higher-order learning and memory processes, and provide a compelling case for exploring antioxidant treatment in individuals with intellectual disability-associated *ALKBH8* variants.

## Discussion

Here, we find that tRNA methyltransferase ALKBH8 promotes redox homeostasis in the nervous system to control synapse development and support associative learning and memory. These findings highlight the critical role of tRNA-modifying enzymes in regulating oxidative stress for nervous system development and function. Notably, the pharmacological rescue of the behavioral phenotype caused by ALKBH8 deficiency suggest antioxidants as a potential therapy for ALKBH8-associated intellectual disability (Figure 6).

**Figure 6.**
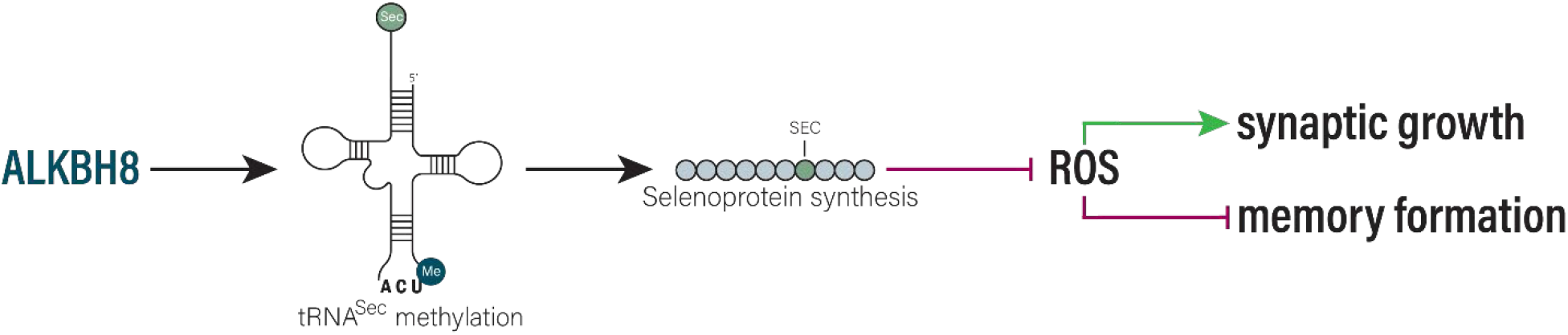
Model of ALKBH8’s role in synapse and memory formation. ALKBH8 methylates tRNA-selenocysteine to promote selenoprotein translation. Selenoproteins control ROS to prevent ectopic synapse formation and promote learning and memory.

ROS act as intracellular signaling molecules in multiple neurodevelopmental processes including differentiation, neurite outgrowth, axon pathfinding, and synaptogenesis (96–99). Our findings demonstrate that while ROS perform key roles during neurodevelopment, their levels must be controlled, with high levels of oxidative stress resulting in ectopic synapse formation. Our findings add to prior studies in *Drosophila* that also observed synaptic overgrowth when ROS levels were elevated (78, 100). At the *Drosophila* NMJ, pathological levels of ROS were found to activate JNK/AP-1 signaling (78), which positively regulates bouton addition (101). ROS signaling is also known to modulate multiple growth factor pathways with roles in synapse development, including the BMP and WNT/Wg pathways (102–108). Investigating whether these pathways also play roles in ROS-induced ectopic synapse formation will be of interest.

ROS are also implicated in synaptic plasticity and homeostasis across species (78, 100, 109, 110). At the *Drosophila* NMJ, Semaphorin-Plexin signaling regulates cytoskeletal dynamics underlying homeostatic modulation of neurotransmitter release through Mical, which uses intrinsic redox potential to oxidize methionine residues in actin and promote depolymerization (111, 112). Activating neurons has also been shown to induce ROS generation (109). ROS are then sensed by DJ-1β, leading to the inhibition of the phosphatase PTEN and activation of phosphatidylinositol 3-kinase (PI3K) signaling, which promotes new synapse formation (109, 113). Thus, signaling downstream of ROS during synapse development and plasticity is diverse and context specific. Our findings suggest that ALKBH8 may act upstream of multiple signaling pathways by regulating selenoprotein synthesis to control ROS levels during synapse development and synaptic plasticity. Notably, ROS can induce *ALKBH8* expression (39, 47), suggesting ALKBH8 may function as part of a feedback mechanism that constrains activity-dependent synapse modification. Intriguingly, a recent study in *Aplysia* demonstrates an increase in mcm^5^s^2^U, a terminal tRNA modification dependent on ALKBH8, during non-associative learning (114).

Our findings suggest that ALKBH8 primarily promotes nervous system development and function by controlling oxidative stress. With their reactive selenocysteine residues, which contain nucleophilic selenium instead of sulfur in cysteine, selenoproteins are potent antioxidant enzymes (68, 115). Selenoprotein synthesis is highly dependent on wobble uridine methylation of tRNA^Sec^ for UGA codon decoding. tRNA^Sec^ is found in two forms in mammals, mcm^5^U catalyzed by ALKBH8 and mcm^5^Um further modified by FTSJ1 (116, 117). Consistent with our findings, the mcm^5^U modification was previously reported to be the sole terminal modification of *Drosophila* tRNA^Sec^ (118). Loss of tRNA^Sec^ modification results in reduced selenoprotein levels in mice as well as flies (44, 47, 49). Across metazoa, small and variable numbers of selenoproteins play key roles in redox homeostasis. Twenty-five selenoproteins have been identified in humans, while only three of these are present in *Drosophila* (70, 119). Importantly, one of these conserved selenoproteins, SelH, is a DNA-binding protein that binds heat shock and stress response elements to regulate the expression of multiple antioxidants, including glutathione transferases (120), suggesting the impact of ALKBH8-dependent tRNA^Sec^ methylation on oxidative stress response extends beyond selenoproteins. Consistently, loss of *Drosophila SPS2*, which is also required for selenoprotein synthesis, results in broad transcriptional downregulation of oxidative stress response genes (121).

Growing links between tRNA methyltransferases and neurodevelopmental disorders underscore the importance of tRNA post-transcriptional modification in the nervous system (16, 18, 27, 122). Variants in *ALKBH8* have now been identified in five families, each resulting in autosomal recessive intellectual disability (58–61). These variants include premature stops in the last exon and missense mutations in residues highly conserved across species, indicating critical roles (123). In all three cases in which they were measured, which included one missense (58) and two nonsense (59) mutations, ALKBH8-dependent tRNA modifications were absent, consistent with complete loss of function. In our studies, methylation of tRNA^Sec^ appears to account for most of ALKBH8’s role in regulating synapse development. However, it is highly likely that loss of ALKBH8-dependent modifications on other tRNAs disrupts translation of additional proteins, as supported by our ability to detect global inhibition of translation in larval motor neurons. The complete suppression of ectopic synaptic growth and learning and memory deficits by genetic rescue or pharmacological treatment with antioxidants, respectively, suggests that these additional proteins may also be linked to redox homeostasis and points to a promising approach for therapeutic intervention.

## Materials and methods

### Drosophila stocks and gene editing

All fly lines were maintained on ready-made Fly Food R medium (LabExpress, Cat# 7003-NV) at 25°C, 60% relative humidity in a 12-hour light-dark cycle incubator (Darwin Chambers). The following fly lines are available at the Bloomington *Drosophila* Stock Center (BDSC): *w^1118^* (BDSC 5905,), P{EPgy2}SPS2[EY00228] (*SPS2*^EY00228^, BDSC 15286), *SPS2* Deficiency (df, BDSC 9635), *UAS-SOD1* (BDSC 24750), *UAS-SOD2* (BDSC 24494), *UAS-catalase* (BDSC 24621), OK371-Gal4 (BDSC 26160; motor-neuronal expression), act5c-Gal4 (BDSC 3954; ubiquitous expression). Full genotypes used in each experiment are described in Table S1. For synaptic bouton quantification experiments, crosses were set up with 10 males and 10 females on Fly Food R medium, placed at 25°C and flipped every 2-3 days to prevent overcrowding.

CRISPR-based homology directed repair was used to generate null and endogenous rescue alleles. Target sites were selected using CRISPR Optimal Target Finder (https://targetfinder.flycrispr.neuro.brown.edu, 124). The following gRNAs were used: 5’ – TTTCCTCAACGTGCATCTCGCGG – 3’ (*ALKBH8*^1-IC^), 5’ – CGTTGAGGAAACACAGCAAAAGG – 3’ (*ALKBH8*^2-IC^) and 5’ – GCAACTTCACCCGGTGTGGGTGG – 3’ (*SPS2*^1^). gRNA and donor plasmids were generated as described in Gratz, Ukken et al., 2014. *Vasa-cas9* embryos were injected with a mixture of a gRNA plasmid (100ng/μL) and a double-stranded DNA donor plasmid (500ng/μL) by BestGene, Inc. and then crossed to *w^1118^* flies after eclosion. Progeny were screened for the presence of DsRed fluorescent expression in the eyes. Endogenously tagged alleles were created using a scarless CRISPR-piggyBac approach (62, 63). A 2xHA (5’– ATGTACCCATACGATGTTCCAGATTACGCTGGATATCCATATGATGTTCCAGATTATGCT – 3’), V5-mCherry (5’ – GTGTCGAAGGGCGAGGAGGATAACATGGCCATCATCAAGGAGTTCATGCGCTTCAAGGTGCACATGGAGGGCTCCGTGAACGGCCACGAGTTCGAGATCGAGGGCGAGGGCGAGGGCCGCCCCTACGAGGGCACCCAGACCGCCAAGCTTAAAGTGACCAAGGGTGGCCCCCTGCCCTTCGCCTGGGACATCCTGTCCCCTCAATTCATGTACGGCTCCAAGGCCTACGTGAAGCACCCCGCCGACATCCCCGACTACTTGAAGCTGTCCTTCCCCGAGGGCTTCAAGTGGGAGCGCGTGATGAACTTCGAGGACGGCGGCGTGGTGACCGTGACCCAGGACTCCTCCCTCCAAGATGGCGAGTTCATTTATAAAGTGAAGCTGCGCGGCACCAACTTCCCCTCCGACGGCCCCGTAATGCAGAAGAAGACCATGGGCTGGGAGGCCTCCTCCGAGCGGATGTACCCCGAGGACGGCGCCCTGAAGGGCGAGATCAAGCAGAGGCTGAAGCTGAAGGACGGCGGCCACTACGACGCTGAGGTCAAGACCACCTACAAGGCCAAGAAGCCCGTGCAGCTGCCCGGCGCCTACAACGTCAACATCAAGTTGGACATCACCTCCCACAACGAGGACTACACCATCGTGGAACAGTACGAACGCGCCGAGGGCCGCCACTCCACCGGCGGCATGGACGAGCTGTACAAG – 3’) or V5 peptide (5’ – GGAAAGCCTATCCCTAACCCTCTCCTCGGTCTCGATTCTACG – 3’) tag flanked by flexible linkers and a visible marker flanked by piggyBac inverted terminal repeat sequences were inserted immediately downstream of the *ALKBH8* and *SPS2* translational start sites, respectively. This insertion generates a null allele due to stop codons in the inverted repeat sequences. piggyBac transposase was subsequently used for scarless removal of the marker cassette. All lines were molecularly verified by PCR and Sanger sequencing.

### Immunostaining, confocal imaging and bouton quantification

Wandering third-instar larvae were dissected in ice cold Ca^2+^-free saline and fixed for 6 minutes in Bouin’s Fixative (Electron Microscopy Sciences, Cat# 15990). Dissections were washed several times with 1X PBS and then permeabilized with 1X PBS containing 0.1% Triton-X (Fisher Scientific, Cat# BP151-100). Dissected larvae were then blocked either for 1 hour at room temperature or overnight at 4°C in 1X PBS containing 0.1% Triton-X and 1% BSA (Sigma-Aldrich, Cat# A7906). Dissections were incubated in primary antibodies overnight at 4°C and secondary antibodies for 2-4 hours at room temperature or overnight at 4°C. Dissected larvae were mounted in Vectashield (Vector Laboratories, Cat# H-1000-10). The following antibodies were used at the indicated concentrations: mouse anti-Bruchpilot (Brp) at 1:100 (DSHB, Cat# nc82, RRID: AB_2314866), Alexa Fluor 488 goat anti-mouse at 1:500 (Invitrogen, Cat# A- 11029, RRID: AB_2534088) and anti-Horseradish Peroxidase (HRP) conjugated to Alexa Fluor 647 at 1:500 (Jackson ImmunoResearch Labs, Cat# 123-605-021, RRID: AB_2338967).

Larval NMJ images were taken on a Nikon A1R HD confocal microscope with a 40x oil immersion objective. Boutons were quantified at the highly stereotyped larval NMJ4 muscle segments, A2-A4. A bouton was defined as a bulbous swelling highlighted by the neuronal membrane marker HRP and co-stained with the active zone marker Brp. Quantification of bouton number was performed masked to genotype/condition using the “multi-point” tool in FIJI/ImageJ (NIH, RRID: SCR_003070).

### Isolation of total tRNA and liquid chromatography-tandem mass spectrometry analysis

Vials containing adult fly heads (mated males and females, three replicates) were flash frozen in liquid nitrogen and stored at −80°C. 25mg of frozen adult heads were homogenized using a plastic pestle in TRIzol reagent. RNA was extracted using an adapted TRIzol extraction protocol (Bogart and Andrews 2006) and resuspended in Rnase-free ddH_2_O. Total RNA (20µg) was nuclease-digested and dephosphorylated with Benzonase, Phosphodiesterase, and Calf Intestinal phosphatase for 3 hours at 37°C to generate individual nucleosides. Enzymes were removed using an Amicon 10,000-Da MWCO spin filter (Millipore, Cat# UFC5010) and purified ribonucleosides were stored at -80°C until further analysis. Ribonucleosides were separated using a Hypersil GOLD™ C18 Selectivity Column (Thermo Scientific, Cat# 25003104630) followed by nucleoside analysis using a Q Exactive Plus Hybrid Quadrupole-Orbitrap (Thermo Scientific) as previously described (39, 125). The mass transitions for each modified nucleoside were validated using modified ribonucleoside standards and known column retention times (126). Mass spectra are quantified by automated integration of peak areas following normalization to the sum of intensity values for the canonical nucleosides A, U, G and C.

### Detection and analysis of protein translation rates via fluorescent non-canonical amino acid tagging (FUNCAT)

To visualize newly synthesized proteins in control and *ALKBH8*^1-IC^ animals, *UAS-dMetRS^L262G^::EGFP* was expressed in larval motor neurons using OK371-Gal4 (66, 67). Following 2-hour egg collections, animals were reared at 25°C on Jazz-Mix *Drosophila* medium (Fisher Scientific, Cat# AS153). Larvae were transferred to medium containing 4mM of non-canonical amino acid azidonorleucine (ANL; Iris Biotech, HAA1625.0025, Cas#: 159610-92- 1net) for 48 hours, followed by larval brain dissection in ice-cold HL3 saline solution (70mM NaCl, 5mM KCl, 20mM MgCl_2_, 10mM NaHCO_3_, 115mM sucrose, 5mM trehalose, 5mM HEPES, pH 7.2) with 0.1mM Ca^2+^ and pre-fixed with 2-3 drops of 4% paraformaldehyde in 1X PBS for 1- minute. Solution was exchanged to 4% paraformaldehyde in 1X PBS and incubated at room temperature for 30-minutes. After fixation, larval brains were washed three times for 10-minutes with PBST (phosphate buffer red saline pH 7.2 + 0.2% v/v Triton X-100) followed by three 10- minute washes with 1X PBS (137mM NaCl, 2.7mM KCl, 4.3mM Na_2_HPO_4_, 1.4mM KH_2_PO_4_, pH 7.8). ANL-labelled proteins were tagged by ’click chemistry’ (Copper-Catalyzed (3+2)-Azide-Alkyne-Cycloaddition Chemistry (CuAAC)) using the fluorescent tag TAMRA. The FUNCAT reaction mix was assembled in a defined sequence of steps. Triazole ligand (1:1000), TAMRA-alkyne tag (1:5000; Sigma Aldrich, Cat# 900932), TCEP solution (1:1000) and CuSO4 solution (1:1000) were added to PBS pH 7.8. After each addition the solution was mixed thoroughly for 10 seconds and at the end for 30 seconds using a high-speed vortex. Larval brains were incubated with 200µl of FUNCAT reaction mix overnight at 4°C on a rotating platform. The next day, the brains were washed 3 times with PBS-Tween and 3 times with PBST for 10-minutes. Finally, larval CNSs were mounted in VectaShield mounting medium (Biozol, Cat# VEC-H-1000- CE) and stored at 4°C until imaging using a Leica SP8 laser scanning confocal microscope. For image acquisition, identical confocal settings were used for all samples of a given experiment. Fluorescence intensities were quantified using FIJI/ImageJ software (NIH). Mean intensity of all cells within one motor neuron cluster from each ventral nerve cord were used as single data points for statistical analysis.

### Protein extraction and Western blot analysis

Protein samples were prepared by dissecting wandering third-instar larvae with intact ventral nerve cords followed by homogenization in Laemmli 2x Sample Buffer (Bio-Rad, Cat# 1610737). Protein lysates were heated at 100°C for 5-minutes. Protein samples were then run on 10% SDS-PAGE at 50V through stacking gel and then at an increased voltage of 100V when samples hit resolving gel. The SDS-PAGE was transferred onto nitrocellulose membranes (Bio-Rad, Cat# 162-0116) at 15V for 30-minutes. To assess efficient transfer, nitrocellulose membranes were quickly exposed to Ponceau-S Staining Solution (Thermo Scientific, Cat# A40000279) before blocking. Transferred membranes were blocked with freshly prepared 5% (weight/volume) BSA in 1X PBS containing 0.1% Tween-20 (Fisher Scientific, Cat# BP33-100) at room temperature for 1 hour and then incubated with mouse anti-V5 (1:5,000; Invitrogen, Cat# R960-25, RRID: AB_2556564) and mouse anti-tubulin (1:4,000; DSHB, Cat# e7-c, RRID: AB_528499) overnight at 4°C. After thorough washing with 1X PBS containing 0.1% Tween-20, blots were incubated with secondary antibody (goat anti-mouse; 1:10,000; Jackson ImmunoResearch Labs, Cat# 115-005-003, RRID: AB_10015289) for 2 hours at room temperature. Blots were thoroughly washed with 1X PBS containing 0.1% Tween-20 and then developed using SuperSignal^TM^ West Femto Maximum Sensitivity Substrate (ThermoFisher Scientific, Cat# 34095) and an Azure c600 Gel Imaging System (Azure Biosystems).

Protein bands from blots were quantified using FIJI/ImageJ (NIH). Briefly, a frame was drawn over the largest protein band in the row using the “rectangle” tool. The defined frame was used to measure all present protein bands. Background measurements were collected using the same defined frame, above and below every protein band as well. The same steps were taken to measure loading control bands. The net protein amount for sample and loading control protein bands were calculated as follows: Mean grey value – Average of background mean grey values. Finally, protein band samples were normalized to relevant loading controls.

### Hydrogen peroxide and paraquat sensitivity assays

Flies were reared on Fly Food R medium at 25°C and flipped every 2-3 days. For H_2_O_2_ exposure, ∼200 F1 progeny were collected per genotype and aged from 3-5 days old. A maximum of 20 flies were placed in each vial and starved for 4 hours on a 1cm by 5cm strip of filter paper saturated with 300µl of milliQ water at 25°C. After starvation, flies were then transferred to new vials containing a 1cm by 5cm strip of filter paper saturated with 300µl of 30% H_2_O_2_ (VWR, Cat# BDH7690-1) and placed at 25°C for 24 hours. For paraquat exposure, 30-40 F1 progeny per genotype were collected and aged from 2-5 days old. Flies were separated into groups of 10 and in vials with filter paper saturated with 200µl of a 5% sucrose containing 50mM paraquat solution and placed at 25°C for 24 hours. After 24 hours of exposure to H_2_O_2_ or paraquat, percent mortality was quantified per genotype per condition. 3 datasets were combined for Paraquat exposure.

### N-acetylcysteine amide (AD4) treatment

AD4 was resuspended in milli-Q water to a final concentration of 100mM and stored in small aliquots at -20°C until needed. B mix Fly Food was made fresh per manufacturer’s instructions (LabExpress, Cat# 3008). Briefly, food mixture was heated to 100°C for 10-minutes to ensure proper incorporation of all ingredients. AD4 or milli-Q water was then added to empty fly vials and mixed with food media (cooled to 65°C) to yield a final concentration of 40μg/ml.

### Taste preference assay and analysis

To establish that differences in PER extension were not due to irregular taste preferences, we conducted a control taste preference assay to ensure that the flies exhibited normal proboscis extension reflex (PER) responses to sucrose and quinine. Pre-mated female flies were starved for 24 hours, mounted on microscope slides, and allowed to acclimate as detailed below. After the rest period, flies were satiated with milli-Q water before applying 500 mM sucrose solution three times to the legs (tarsi). After washing the tarsi with water, quinine solution was applied three times to the tarsi by an experimenter masked to genotype and treatment. We observed the number of PER extensions for both solutions. Solutions were presented to the flies using a 1mL syringe. If the fly exhibited a proboscis extension or no extension, they were given a score of 1 or 0, respectively. The performance index was calculated as the average number of PER displayed out of the 3 presentations multiplied by 100 of either the sucrose or quinine solutions. A high-performance index indicates that the solution presented was appetitive, whereas a low performance index reveals an aversion to the applied solution.

### Aversion memory assay and analysis

The following protocol was adapted from Masek et al. 2010, 2015 and Kirkhart and Scott, 2015. 30-40 mated female flies were starved for 24 hours at 25°C in vials containing 1cm by 6cm strips of filter paper saturated with 300μl of DI water. After the starvation period, the flies were anesthetized using CO_2_ gas and mounted to microscope slides using clear nail polish (Electron Microscopy Sciences, Cat# 72180) applied to the wing base and dorsal side of the thorax by an experimenter masked to genotype and treatment. Ten flies were mounted across two rows on each slide. 500mM sucrose (Sigma-Aldrich, Cat# S7903) and 50mM quinine dihydrochloride (Sigma-Aldrich Cat# Q1125) solutions were freshly prepared. After mounting, the slides were kept upright in a dark, moist chamber for 2.5 hours, allowing the flies to acclimate and recover from CO_2_ gas exposure. After the rest period, flies were satiated with milli-Q water before applying the 500mM sucrose solution three times to the legs (tarsi). Solutions were presented to flies using a 1mL syringe. The number of proboscis extension reflexes (PER) that each fly displayed were recorded; any flies that did not show PER to all three applications of sucrose in the pretest phase were excluded from later analysis since these flies exhibit an abnormal taste response. After the pre-test phase, tarsi were washed with water. Flies were then trained by applying sucrose to their tarsi coupled with immediate presentation of quinine to their extended proboscis three times with 2-minute rest periods between each bout of training, during which tarsi were washed with water. After the third training period, flies were allowed to rest for 5- minutes before applying sucrose to the tarsi three times and recording PER suppression response. To calculate the PER suppression response, we computed a PER suppression index for each fly of 1 minus the fraction of PER displayed in the testing period out of 3 possible responses multiplied by 100. Thus, a high PER suppression percentage indicates that the fly learned to associate quinine with a sucrose cue.

### Experimental design and statistical analyses

All analyses were performed masked to genotype and/or treatment. Statistical analyses were conducted in GraphPad Prism 9 (RRID: SCR_002798). The D’Agostino-Pearson omnibus test was used to assess normality. Single comparisons of normally distributed datasets were analyzed by Student’s t test. Welch’s correction was used in cases of unequal variance. The Mann-Whitney U test was used for single comparisons of non-normally distributed data. For multiple comparisons of normally distributed data with equal variance or unequal variance, One-way ANOVA followed by Tukey’s test or Dunnett’s T3 multiple comparisons test was performed, respectively. Multiple comparisons of non-normally distributed data were performed using the Kruskal-Wallis test followed by Dunn’s multiple comparisons test. Significance is reported as values less than 0.05, 0.01, 0.001, and 0.0001 represented by one, two, three, or four stars, respectively. Unless otherwise noted, significance refers to the indicated genotype compared to control. Reported values are mean ± SEM. Sample sizes and statistics are provided in Table S1.

### Data, materials, and software availability

Source data for all figures will be made available through the Harvard Dataverse repository.

## Acknowledgments

We thank the Developmental Studies Hybridoma Bank and Bloomington Drosophila Stock Center for providing antibodies and fly stocks, and Kevin Welle and the Mass Spectrometry Resource Lab at the University of Rochester. We are grateful to the O’Connor-Giles, Fu, and Storkebaum labs, Karla Kaun, and Kristi Wharton for critical feedback on this work and manuscript. This work was supported by a grant from the NIH National Institute of Neurological Disorders and Stroke (NINDS) to K.M.O.-G. and D.F. (R01NS117068). K.R.M. was supported by NINDS F99NS129128, J.M.L. and D.F. by NSF CAREER Award 1552126 and NIH R01GR532955, and J.L.D. through the Brown University Predoctoral Training Program in Trans-disciplinary Pharmacological Sciences (T32 GM077995). E.S. is supported by an ERC consolidator grant (ERC-2017-COG 770244), and funding from the Radala Foundation, ’Stichting ALS Nederland’, AFM-Telethon, ARSLA, the ‘Prinses Beatrix Spierfonds’ (W.OR22- 03), the Muscular Dystrophy Association (MDA 946876) and an NWO Open Competition ENW-M grant.

---

**Figure S1.**
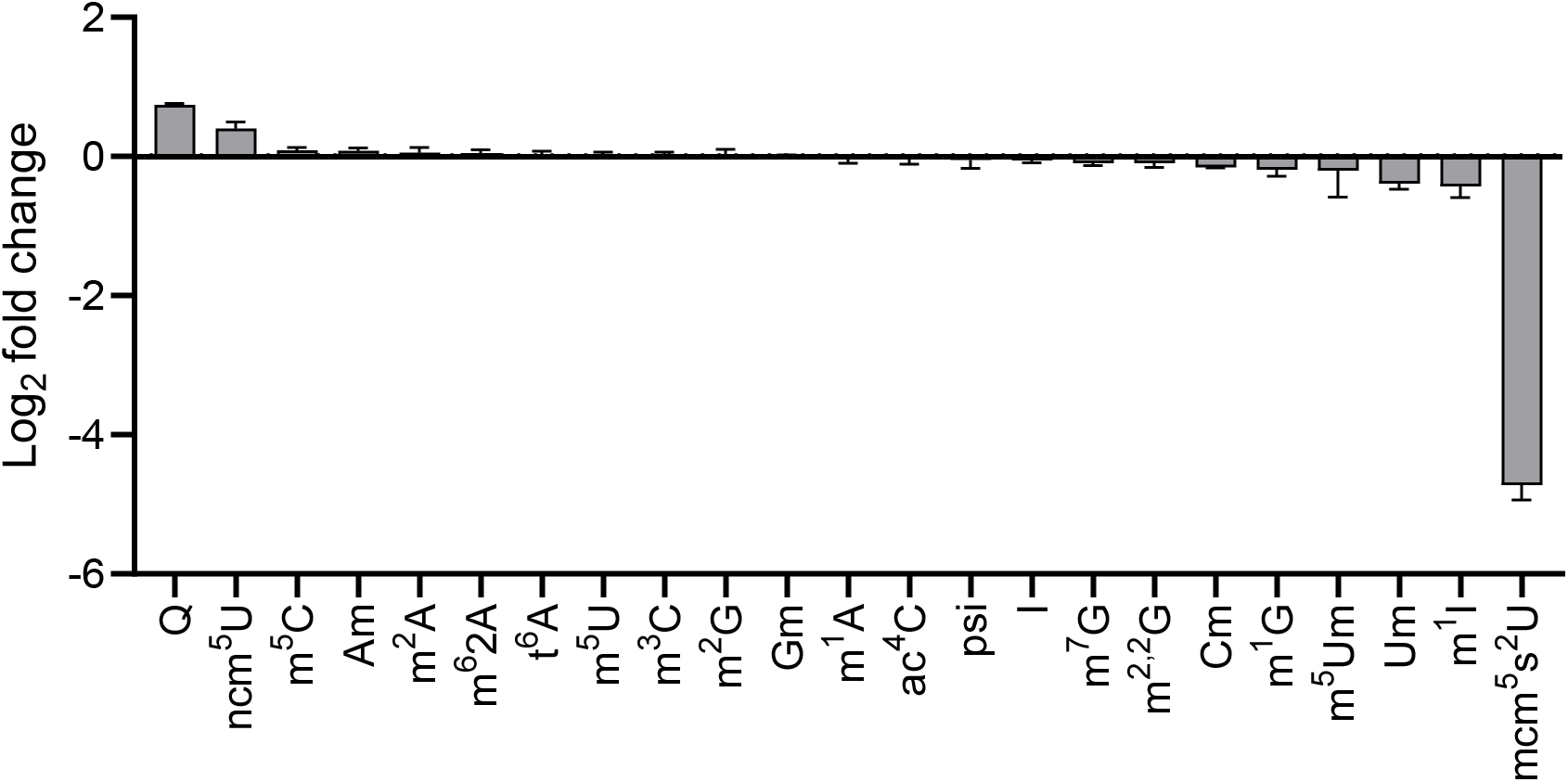
Other tRNA modifications are not impacted in *ALKBH8* null animals. Log_2_ fold change in levels of the indicated tRNA modifications in *ALKBH8*^1-IC^ null heads relative to control. 3 biological replicates per genotype. Data is represented as mean ± SEM.

**Figure S2.**
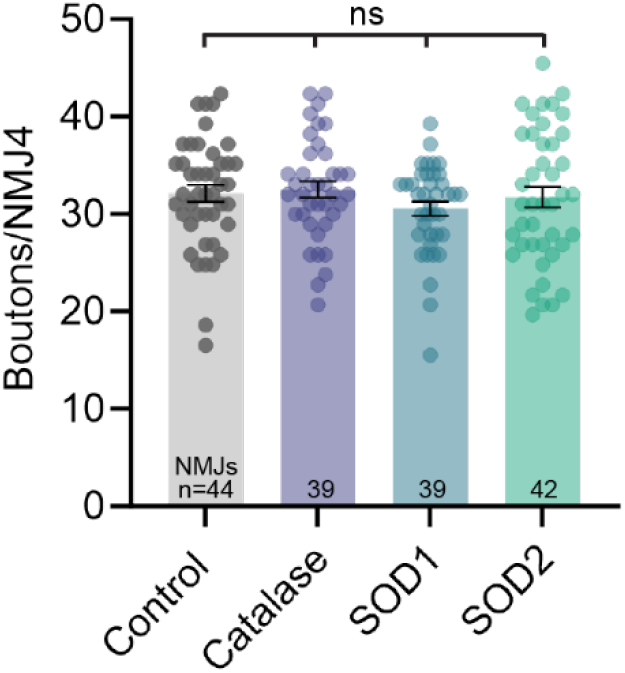
Overexpression of antioxidant enzymes does not impact synaptic growth. Ubiquitous overexpression of antioxidant enzymes in a control background does not affect synapse formation. Data points represent individual NMJs from 39-44 animals with n indicated in graph. Kruskal-Wallis test followed by Dunn’s multiple comparisons test. Data is represented as mean ± SEM. *ns* = not significant.

**Figure S3.**
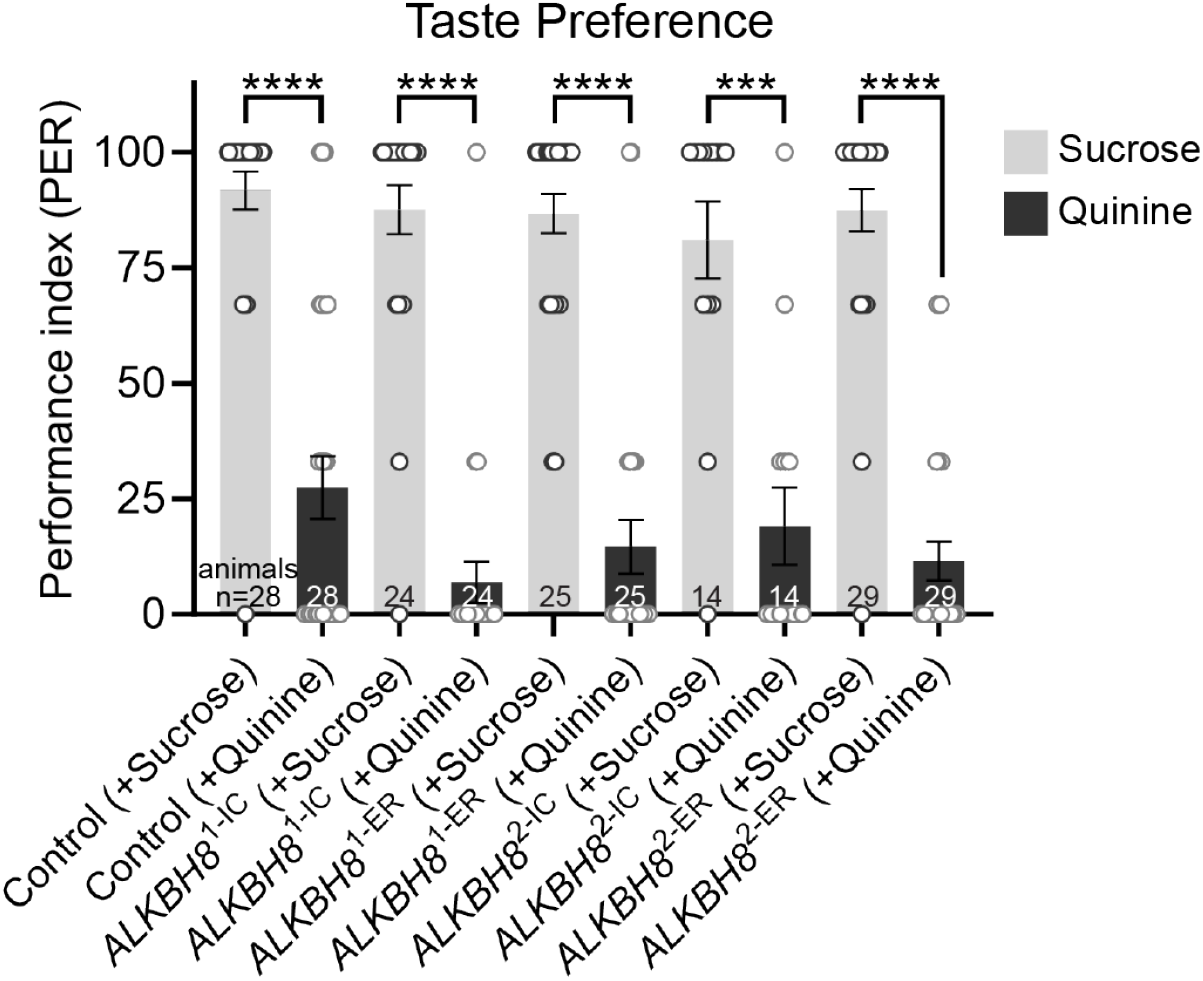
*ALKBH8* null mutants exhibit normal taste preference. All genotypes exhibit expected PER responses when presented with either sucrose or quinine. Data points represent individual animals with n indicated in graphs. Mann-Whitney U test. Data is represented as mean ± SEM. \*\*\*\**p* < 0.0001, \*\*\**p* = 0.0002.

**Table S1.**
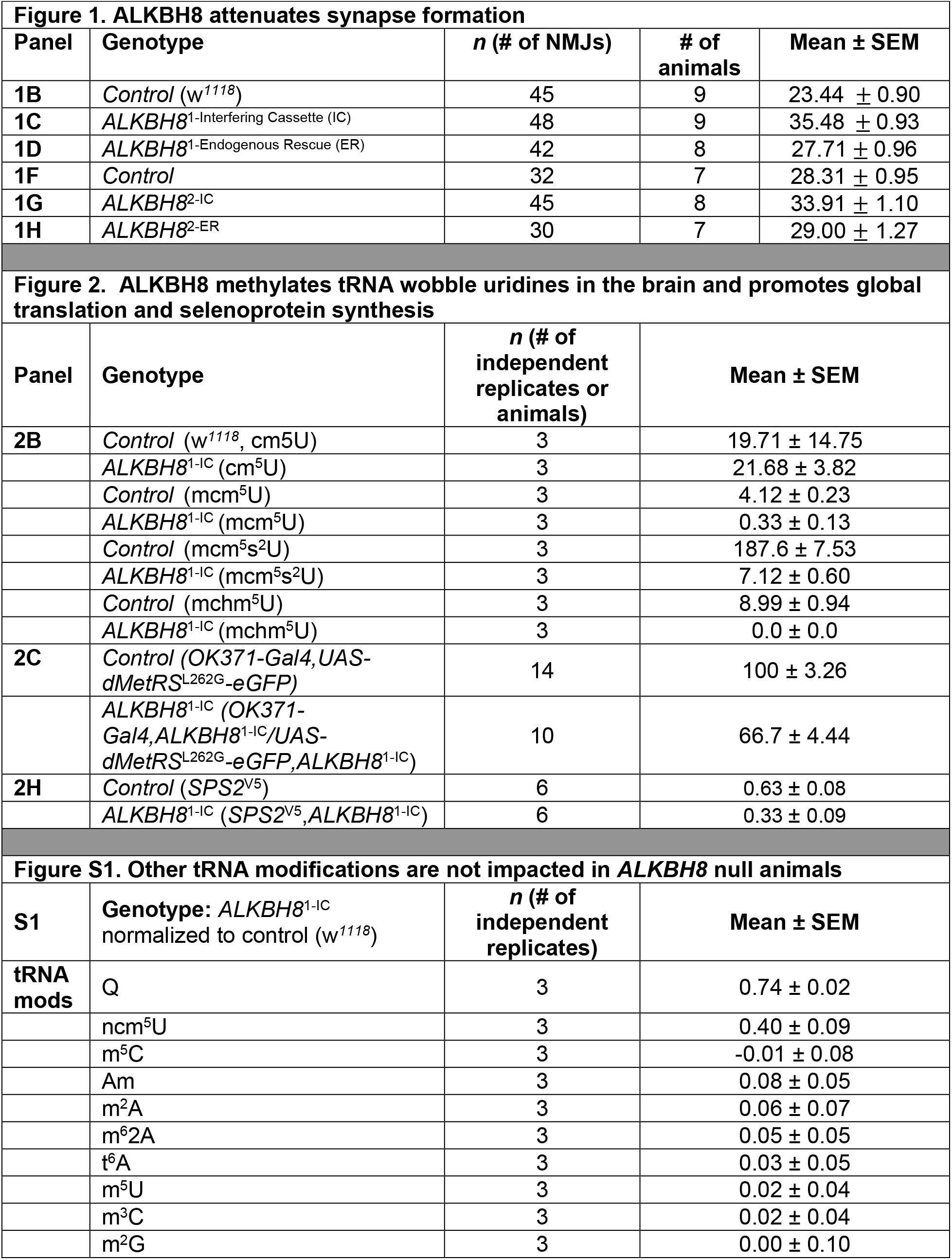
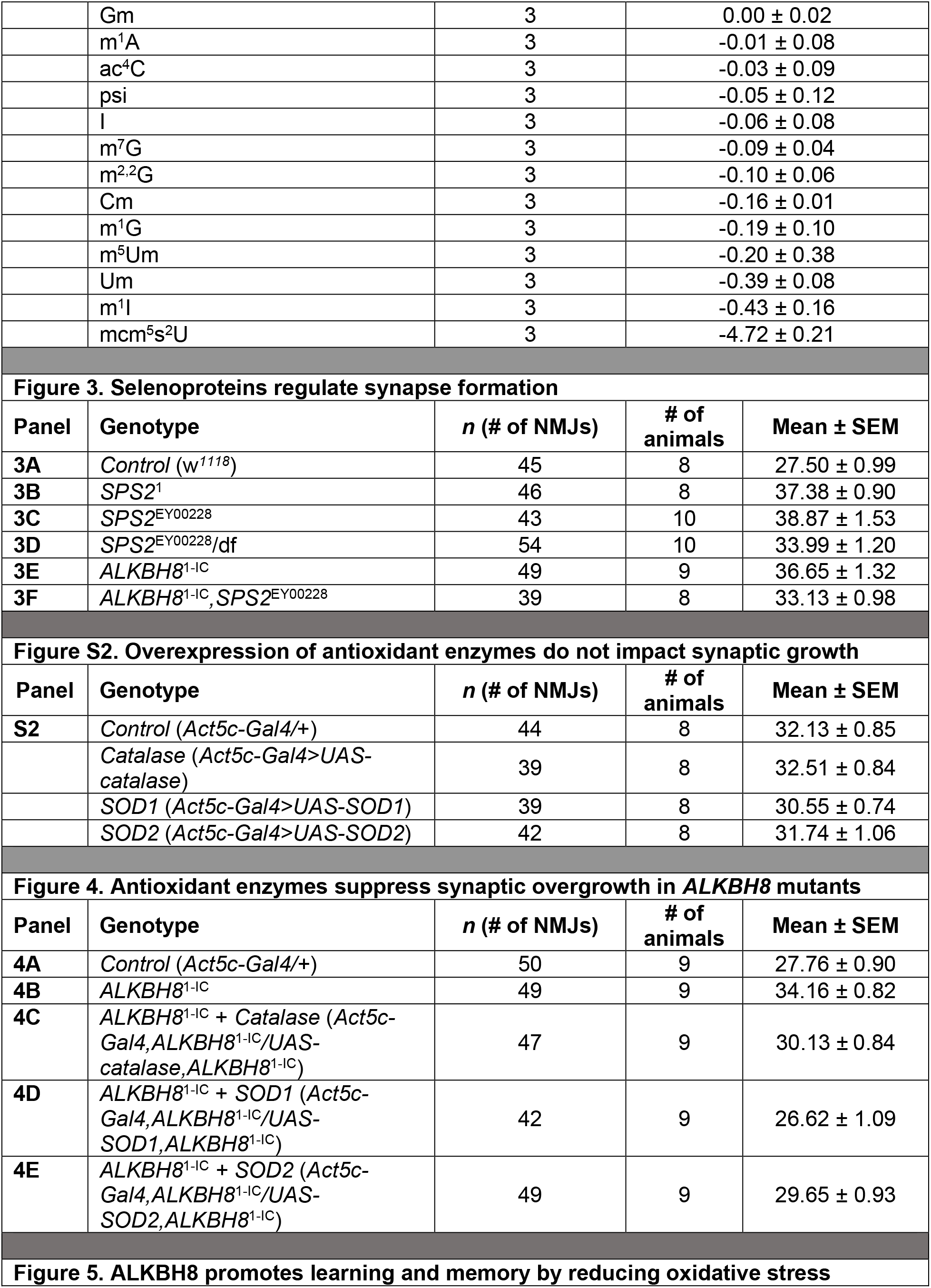
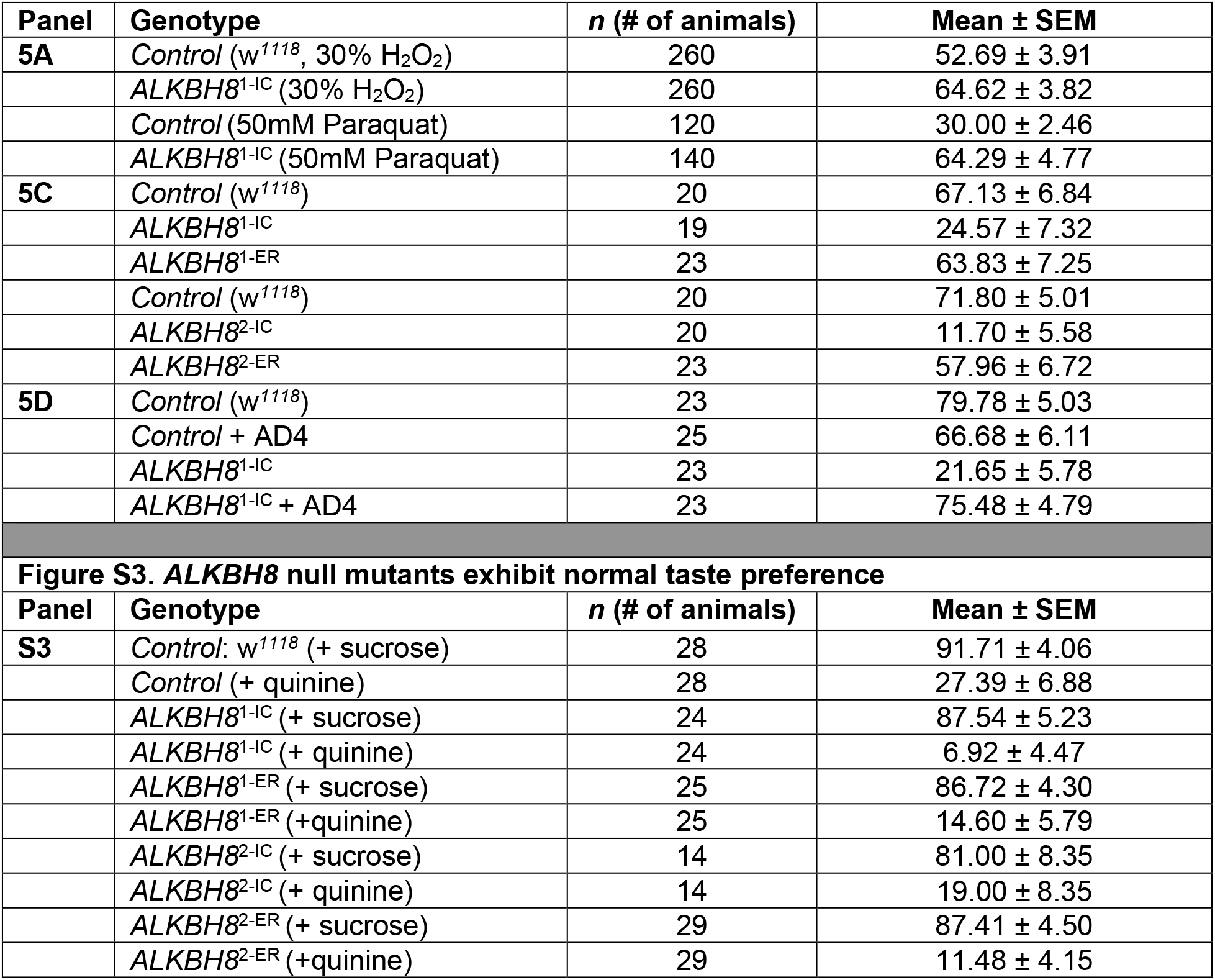
Absolute values for all data. Full genotypes and/or treatments, sample size, and mean ± SEM are provided for each figure below.

## Notes

### Competing Interest Statement

The authors have declared no competing interest.

## References

1. S. M. Carpanini, T. M. Wishart, T. H. Gillingwater, J. C. Manson, K. M. Summers, Analysis of gene expression in the nervous system identifies key genes and novel candidates for health and disease. neurogenetics 18, 81–95 (2017).

2. S. W. Flavell, M. E. Greenberg, Expression and Plasticity of the Nervous System. Annu Rev Neurosci, 563–590 (2008).

3. M. Gamarra, A. De La Cruz, M. Blanco-Urrejola, J. Baleriola, Local Translation in Nervous System Pathologies. Front. Integr. Neurosci. 15, 689208 (2021).

4. K. Gapp, B. T. Woldemichael, J. Bohacek, I. M. Mansuy, Epigenetic regulation in neurodevelopment and neurodegenerative diseases. Neuroscience 264, 99–111 (2014).

5. C. E. Holt, E. M. Schuman, The central dogma decentralized: New perspectives on RNA function and local translation in neurons. Neuron 80, 648–657 (2013).

6. L. Ooi, I. C. Wood, Regulation of gene expression in the nervous system. Biochem. J. 414, 327–341 (2008).

7. Z. Qiu, A. Ghosh, A Brief History of Neuronal Gene Expression: Regulatory Mechanisms and Cellular Consequences. Neuron 60, 449–455 (2008).

8. R. D. Salinas, D. R. Connolly, H. Song, Invited Review: Epigenetics in neurodevelopment. Neuropathol. Appl. Neurobiol. 46, 6–27 (2020).

9. W. S. Sossin, M. Costa-Mattioli, Translational Control in the Brain in Health and Disease. Cold Spring Harb. Perspect. Biol. 11, a032912 (2019).

10. Y. Chen, Y. Wang, Z. Modrusan, M. Sheng, J. S. Kaminker, Regulation of Neuronal Gene Expression and Survival by Basal NMDA Receptor Activity: A Role for Histone Deacetylase 4. J. Neurosci. 34, 15327–15339 (2014).

11. D. W. Hwang, et al., Chromatin-mediated translational control is essential for neural cell fate specification. Life Sci. Alliance 1, 1–13 (2018).

12. K. Keleman, S. Krüttner, M. Alenius, B. J. Dickson, Function of the Drosophila CPEB protein Orb2 in long-term courtship memory. Nat. Neurosci. 10, 1587–1593 (2007).

13. M. D. Berg, C. J. Brandl, Transfer RNAs: diversity in form and function. RNA Biol. 18, 316–339 (2021).

14. J. M. Ogle, et al., Recognition of Cognate Transfer RNA by the 30S Ribosomal Subunit Published by : American Association for the Advancement of Science Stable URL : http://www.jstor.org/stable/3083592 Recognition of Cognate Transfer RNA by the 30S Ribosomal Subunit. 292, 897–902 (2001).

15. C. Andreassi, H. Crerar, A. Riccio, Post-transcriptional processing of mRNA in neurons: The vestiges of the RNA world drive transcriptome diversity. Front. Mol. Neurosci. 11, 1– 16 (2018).

16. R. W. Burgess, E. Storkebaum, tRNA Dysregulation in Neurodevelopmental and Neurodegenerative Diseases. Annu. Rev. Cell Dev. Biol. 39, annurev-cellbio-021623- 124009 (2023).

17. M. Pereira, et al., Impact of tRNA modifications and tRNA-modifying enzymes on proteostasis and human disease. Int. J. Mol. Sci. 19 (2018).

18. J. Ramos, D. Fu, The emerging impact of tRNA modifications in the brain and nervous system. Biochim. Biophys. Acta - Gene Regul. Mech. 1862, 412–428 (2019).

19. B. El Yacoubi, M. Bailly, V. de Crécy-Lagard, Biosynthesis and Function of Posttranscriptional Modifications of Transfer RNAs. Annu. Rev. Genet. 46, 69–95 (2012).

20. H. Hori, Methylated nucleosides in tRNA and tRNA methyltransferases. Front. Genet. 5, 1–26 (2014).

21. J. E. Jackman, J. D. Alfonzo, Transfer RNA modifications: Nature’s combinatorial chemistry playground. Wiley Interdiscip. Rev. RNA 4, 35–48 (2013).

22. S. Kirchner, Z. Ignatova, Emerging roles of tRNA in adaptive translation, signalling dynamics and disease. Nat. Rev. Genet. 16, 98–112 (2015).

23. P. O’Donoghue, J. Ling, D. Söll, Transfer RNA function and evolution. RNA Biol. 15, 423– 426 (2018).

24. I. A. Roundtree, M. E. Evans, T. Pan, C. He, Dynamic RNA Modifications in Gene Expression Regulation. Cell 169, 1187–1200 (2017).

25. W. E. Swinehart, J. E. Jackman, Diversity in mechanism and function of tRNA methyltransferases. RNA Biol. 12, 398–411 (2015).

26. W. Zhang, M. Foo, A. M. Eren, T. Pan, tRNA modification dynamics from individual organisms to metaepitranscriptomics of microbiomes. Mol. Cell 82, 891–906 (2022).

27. A. Bednářová, et al., Lost in translation: Defects in transfer RNA modifications and neurological disorders. Front. Mol. Neurosci. 10, 1–8 (2017).

28. W. Cui, et al., tRNA Modifications and Modifying Enzymes in Disease, the Potential Therapeutic Targets. Int. J. Biol. Sci. 19, 1146–1162 (2023).

29. A. E. Schaffer, O. Pinkard, J. M. Coller, TRNA Metabolism and Neurodevelopmental Disorders (2019).

30. T. Suzuki, The expanding world of tRNA modifications and their disease relevance. Nat. Rev. Mol. Cell Biol. 22, 375–392 (2021).

31. J.-B. Zhou, E.-D. Wang, X.-L. Zhou, Modifications of the human tRNA anticodon loop and their associations with genetic diseases. Cell. Mol. Life Sci. 78, 7087–7105 (2021).

32. A. Rozov, et al., Novel base-pairing interactions at the tRNA wobble position crucial for accurate reading of the genetic code. Nat. Commun. 7, 1–10 (2016).

33. R. Schaffrath, S. A. Leidel, Wobble uridine modifications – a reason to live, a reason to die?! 14, 1209–1222 (2017).

34. H. R. Kalhor, S. Clarke, Novel Methyltransferase for Modified Uridine Residues at the Wobble Position of tRNA. Mol. Cell. Biol. 23, 9283–9292 (2003).

35. U. Begley, et al., Trm9-Catalyzed tRNA Modifications Link Translation to the DNA Damage Response. Mol. Cell 28, 860–870 (2007).

36. C. T. Y. Chan, et al., A quantitative systems approach reveals dynamic control of tRNA modifications during cellular stress. PLoS Genet. 6, 1–9 (2010).

37. W. Deng, et al., Trm9-Catalyzed tRNA Modifications Regulate Global Protein Expression by Codon-Biased Translation. PLoS Genet. 11, 1–23 (2015).

38. A. Patil, et al., Translational infidelity-induced protein stress results from a deficiency in Trm9-catalyzed tRNA modifications. RNA Biol. 9, 990–1001 (2012).

39. D. Fu, et al., Human AlkB Homolog ABH8 Is a tRNA Methyltransferase Required for Wobble Uridine Modification and DNA Damage Survival. Mol. Cell. Biol. 30, 2449–2459 (2010).

40. Y. Fu, et al., The AlkB Domain of Mammalian ABH8 Catalyzes Hydroxylation of 5- Methoxycarbonylmethyluridine at the Wobble Position of tRNA. Angew. Chem. Int. Ed. 49, 8885–8888 (2010).

41. C. A. Hogan, et al., Expanded TRNA methyltransferase family member TRMT9B regulates synaptic growth and function. EMBO Rep., e56808 (2023).

42. J. Kollárová, M. Kostrouchová, A. Benda, M. Kostrouchová, ALKB-8, a 2-Oxoglutarate-Dependent Dioxygenase and S-Adenosine Methionine-Dependent Methyltransferase Modulates Metabolic Events Linked to Lysosome-Related Organelles and Aging in C. elegans. Folia Biol (Praha) 64, 46–58 (2018).

43. V. Leihne, et al., Roles of Trm9-and ALKBH8-like proteins in the formation of modified wobble uridines in Arabidopsis tRNA. Nucleic Acids Res. 39, 7688–7701 (2011).

44. L. Songe-Moller, et al., Mammalian ALKBH8 Possesses tRNA Methyltransferase Activity Required for the Biogenesis of Multiple Wobble Uridine Modifications Implicated in Translational Decoding. Mol. Cell. Biol. 30, 1814–1827 (2010).

45. E. Van Den Born, et al., ALKBH8-mediated formation of a novel diastereomeric pair of wobble nucleosides in mammalian tRNA. Nat. Commun. 2 (2011).

46. I. Cavallin, et al., HITS-CLIP analysis of human ALKBH8 reveals interactions with fully processed substrate tRNAs and with specific noncoding RNAs. RNA, rna.079421.122 (2022).

47. L. Endres, et al., Alkbh8 regulates selenocysteine-protein expression to protect against reactive oxygen species damage. PLoS ONE 10, 1–23 (2015).

48. S. Evke, Q. Lin, J. A. Melendez, T. J. Begley, Epitranscriptomic Reprogramming Is Required to Prevent Stress and Damage from Acetaminophen. Genes 13, 421 (2022).

49. M. Y. Lee, A. Leonardi, T. J. Begley, J. A. Melendez, Loss of epitranscriptomic control of selenocysteine utilization engages senescence and mitochondrial reprogramming☆. Redox Biol. 28, 101375 (2020).

50. M. Y. Lee, et al., Selenoproteins and the senescence-associated epitranscriptome. Exp. Biol. Med. 247, 2090–2102 (2022).

51. A. Leonardi, S. Evke, M. Lee, J. A. Melendez, T. J. Begley, Epitranscriptomic systems regulate the translation of reactive oxygen species detoxifying and disease linked selenoproteins. Free Radic. Biol. Med. 143, 573–593 (2019).

52. A. Leonardi, et al., The epitranscriptomic writer ALKBH8 drives tolerance and protects mouse lungs from the environmental pollutant naphthalene. Epigenetics 00, 1–18 (2020).

53. O. Agamy, et al., Mutations disrupting selenocysteine formation cause progressive cerebello-cerebral atrophy. Am. J. Hum. Genet. 87, 538–544 (2010).

54. A. K. Anttonen, et al., Selenoprotein biosynthesis defect causes progressive encephalopathy with elevated lactate. Neurology 85, 306–315 (2015).

55. S. Behl, S. Mehta, M. K. Pandey, The role of selenoproteins in neurodevelopment and neurological function: Implications in autism spectrum disorder. Front. Mol. Neurosci. 16, 1130922 (2023).

56. E. Schoenmakers, K. Chatterjee, Human Genetic Disorders Resulting in Systemic Selenoprotein Deficiency. Int. J. Mol. Sci. 22, 12927 (2021).

57. U. Schweizer, M. Fabiano, Selenoproteins in brain development and function. Free Radic. Biol. Med. 190, 105–115 (2022).

58. S. Maddirevula, et al., Insight into ALKBH8-related intellectual developmental disability based on the first pathogenic missense variant. Hum. Genet., 1–7 (2021).

59. D. Monies, C. B. Vågbø, M. Al-Owain, S. Alhomaidi, F. S. Alkuraya, Recessive Truncating Mutations in ALKBH8 Cause Intellectual Disability and Severe Impairment of Wobble Uridine Modification. Am. J. Hum. Genet. 104, 1202–1209 (2019).

60. A. K. Saad, et al., Neurodevelopmental disorder in an Egyptian family with a biallelic ALKBH8 variant. Am. J. Med. Genet. A. 185, 1288–1293 (2021).

61. A. Waqas, et al., Case Report: Biallelic Variant in the tRNA Methyltransferase Domain of the AlkB Homolog 8 Causes Syndromic Intellectual Disability. Front. Genet. 13, 878274 (2022).

62. J. J. Bruckner, et al., Fife organizes synaptic vesicles and calcium channels for high-probability neurotransmitter release. J. Cell Biol. 216, 231–246 (2017).

63. S. J. Gratz, et al., Endogenous tagging reveals differential regulation of Ca 2+ channels at single AZs during presynaptic homeostatic potentiation and depression. J. Neurosci., 3068–18 (2019).

64. C. A. Collins, A. DiAntonio, Synaptic development: insights from Drosophila. Curr. Opin. Neurobiol. 17, 35–42 (2007).

65. W. M. Cai, et al., “A Platform for Discovery and Quantification of Modified Ribonucleosides in RNA” in Methods in Enzymology, (Elsevier, 2015), pp. 29–71.

66. I. Erdmann, et al., Cell-selective labelling of proteomes in Drosophila melanogaster. Nat. Commun. 6, 7521 (2015).

67. S. Niehues, et al., Impaired protein translation in Drosophila models for Charcot–Marie– Tooth neuropathy caused by mutant tRNA synthetases. Nat. Commun. 6, 7520 (2015).

68. V. M. Labunskyy, D. L. Hatfield, V. N. Gladyshev, Selenoproteins: Molecular Pathways and Physiological Roles. Physiol. Rev. 94, 739–777 (2014).

69. S. Castellano, et al., In silico identification of novel selenoproteins in the Drosophila melanogaster genome. EMBO Rep. 2, 697–702 (2001).

70. F. J. Martin-Romero, et al., Selenium metabolism in Drosophila. Selenoproteins, selenoprotein mRNA expression, fertility, and mortality. J. Biol. Chem. 276, 29798–29804 (2001).

71. U. Schweizer, N. Fradejas-Villar, Why 21? the significance of selenoproteins for human health revealed by inborn errors of metabolism. FASEB J. 30, 3669–3681 (2016).

72. M. Hirosawa-Takamori, H. Jäckle, G. Vorbrüggen, The class 2 selenophosphate synthetase gene of Drosophila contains a functional mammalian-type SECIS. EMBO Rep. 1, 441–446 (2000).

73. J. S. Jin, A DNA replication-related element downstream from the initiation site of Drosophila selenophosphate synthetase 2 gene is essential for its transcription. Nucleic Acids Res. 32, 2482–2493 (2004).

74. D. L. Hatfield, B. A. Carlson, X. Xu, H. Mix,, V. N. Gladyshev, “Selenocysteine Incorporation Machinery and the Role of Selenoproteins in Development and Health” in Progress in Nucleic Acid Research and Molecular Biology, (Elsevier, 2006), pp. 97–142.

75. H. J. Bellen, et al., The BDGP gene disruption project: Single transposon insertions associated with 40% of Drosophila genes. Genetics 167, 761–781 (2004).

76. A. Nandi, L.-J. Yan, C. K. Jana, N. Das, Role of Catalase in Oxidative Stress-and Age-Associated Degenerative Diseases. Oxid. Med. Cell. Longev. (2019) 10.1155/2019/9613090.

77. Y. Wang, R. Branicky, A. Noë, S. Hekimi, Superoxide dismutases: Dual roles in controlling ROS damage and regulating ROS signaling. J. Cell Biol. 217, 1915–1928 (2018).

78. V. J. Milton, et al., Oxidative stress induces overgrowth of the Drosophila neuromuscular junction. Proc. Natl. Acad. Sci. U. S. A. 108, 17521–17526 (2011).

79. S. L. DeAngelo, B. Győrffy, M. Koutmos, Y. M. Shah, Selenoproteins and tRNA-Sec: regulators of cancer redox homeostasis. Trends Cancer, S2405803323001632 (2023).

80. V. A. Shchedrina, et al., Analyses of fruit flies that do not express selenoproteins or express the mouse selenoprotein, methionine sulfoxide reductase B1,reveal a role of selenoproteins in stress resistance. J. Biol. Chem. 286, 29449–29461 (2011).

81. Y. Zhang, et al., Role of Selenoproteins in Redox Regulation of Signaling and the Antioxidant System: A Review. Antioxidants 9, 383 (2020).

82. M. Baek, W. Jang, C. Kim, Dual Oxidase, a Hydrogen-Peroxide-Producing Enzyme, Regulates Neuronal Oxidative Damage and Animal Lifespan in Drosophila melanogaster. Cells 11, 2059 (2022).

83. D. Grover, et al., Hydrogen Peroxide Stimulates Activity and Alters Behavior in Drosophila melanogaster. PLoS ONE 4, e7580 (2009).

84. H. Sies, Hydrogen peroxide as a central redox signaling molecule in physiological oxidative stress: Oxidative eustress. Redox Biol. 11, 613–619 (2017).

85. Y. Zou, et al., Cordyceps sinensis oral liquid prolongs the lifespan of the fruit fly, Drosophila melanogaster, by inhibiting oxidative stress. Int. J. Mol. Med. 36, 939–946 (2015).

86. E. Bonilla, S. Medina-Leendertz, V. Villalobos, L. Molero, A. Bohórquez, Paraquat-induced Oxidative Stress in Drosophila melanogaster: Effects of Melatonin, Glutathione, Serotonin, Minocycline, Lipoic Acid and Ascorbic Acid. Neurochem. Res. 31, 1425–1432 (2006).

87. R. Hosamani, Muralidhara, ACUTE EXPOSURE OF *Drosophila melanogaster* TO PARAQUAT CAUSES OXIDATIVE STRESS AND MITOCHONDRIAL DYSFUNCTION: Acute Paraquat-Induced Oxidative Stress in *Drosophila*. Arch. Insect Biochem. Physiol. 83, 25–40 (2013).

88. U. Maitra, M. N. Scaglione, S. Chtarbanova, J. M. O’Donnell, Innate immune responses to paraquat exposure in a Drosophila model of Parkinson’s disease. Sci. Rep. 9, 12714 (2019).

89. T. Z. Rzezniczak, L. A. Douglas, J. H. Watterson, T. J. S. Merritt, Paraquat administration in Drosophila for use in metabolic studies of oxidative stress. Anal. Biochem. 419, 345– 347 (2011).

90. F. Gámiz, Taste learning and memory: a window on the study of brain aging. Front. Syst. Neurosci. 5 (2011).

91. P. Masek, K. Scott, Limited taste discrimination in *Drosophila*. Proc. Natl. Acad. Sci. 107, 14833–14838 (2010).

92. A. C. Keene, P. Masek, Optogenetic induction of aversive taste memory. Neuroscience 222, 173–180 (2012).

93. C. Kirkhart, K. Scott, Gustatory Learning and Processing in the *Drosophila* Mushroom Bodies. J. Neurosci. 35, 5950–5958 (2015).

94. P. Masek, K. Worden, Y. Aso, G. M. Rubin, A. C. Keene, A Dopamine-Modulated Neural Circuit Regulating Aversive Taste Memory in Drosophila. Curr. Biol. 25, 1535–1541 (2015).

95. P. Masek, A. C. Keene, Gustatory processing and taste memory in *Drosophila*. J. Neurogenet. 30, 112–121 (2016).

96. K. Biswas, K. Alexander, M. M. Francis, Reactive Oxygen Species: Angels and Demons in the Life of a Neuron. NeuroSci 3, 130–145 (2022).

97. V. J. Milton, S. T. Sweeney, Oxidative stress in synapse development and function. Dev. Neurobiol. 72, 100–110 (2012).

98. M. C. W. Oswald, N. Garnham, S. T. Sweeney, M. Landgraf, Regulation of neuronal development and function by ROS. FEBS Lett. 592, 679–691 (2018).

99. C. Wilson, E. Muñoz-Palma, C. González-Billault, From birth to death: A role for reactive oxygen species in neuronal development. Semin. Cell Dev. Biol. 80, 43–49 (2018).

100. J.-J. Peng, et al., A circuit-dependent ROS feedback loop mediates glutamate excitotoxicity to sculpt the Drosophila motor system. eLife 8, e47372 (2019).

101. S. Sanyal, D. J. Sandstrom, C. A. Hoeffer, M. Ramaswami, AP-1 functions upstream of CREB to control synaptic plasticity in Drosophila. Nature 416, 870–874 (2002).

102. H. Aberle, et al., wishful thinking Encodes a BMP Type II Receptor that Regulates Synaptic Growth in Drosophila. Neuron 33, 545–558 (2002).

103. S. Chatterjee, P. C. Sil, ROS-Influenced Regulatory Cross-Talk With Wnt Signaling Pathway During Perinatal Development. Front. Mol. Biosci. 9, 889719 (2022).

104. G. Marqués, et al., The Drosophila BMP Type II Receptor Wishful Thinking Regulates Neuromuscular Synapse Morphology and Function. Neuron 33, 529–543 (2002).

105. M. Packard, et al., The Drosophila Wnt, Wingless, Provides an Essential Signal for Pre-and Postsynaptic Differentiation. Cell 111, 319–330 (2002).

106. V. Y. Poon, S. Choi, M. Park, Growth factors in synaptic function. Front. Synaptic Neurosci. 5 (2013).

107. P. C. Salinas, Wnt Signaling in the Vertebrate Central Nervous System: From Axon Guidance to Synaptic Function. Cold Spring Harb. Perspect. Biol. 4, a008003–a008003 (2012).

108. C. Sánchez-de-Diego, J. A. Valer, C. Pimenta-Lopes, J. L. Rosa, F. Ventura, Interplay between BMPs and Reactive Oxygen Species in Cell Signaling and Pathology. Biomolecules 9, 534 (2019).

109. M. C. W. Oswald, et al., Reactive oxygen species regulate activity-dependent neuronal plasticity in Drosophila. eLife 7 (2018).

110. C. Ugbode, et al., JNK signalling regulates antioxidant responses in neurons. Redox Biol. 37, 101712 (2020).

111. B. O. Orr, R. D. Fetter, G. W. Davis, Retrograde semaphorin–plexin signalling drives homeostatic synaptic plasticity. Nature 550, 109–113 (2017).

112. H. Wu, H. G. Yesilyurt, J. Yoon, J. R. Terman, The MICALs are a Family of F-actin Dismantling Oxidoreductases Conserved from Drosophila to Humans. Sci. Rep. 8, 937 (2018).

113. A. Martín-Peña, et al., Age-Independent Synaptogenesis by Phosphoinositide 3 Kinase. J. Neurosci. 26, 10199–10208 (2006).

114. K. D. Clark, C. Lee, R. Gillette, J. V. Sweedler, Characterization of Neuronal RNA Modifications during Non-associative Learning in *Aplysia* Reveals Key Roles for tRNAs in Behavioral Sensitization. ACS Cent. Sci. 7, 1183–1190 (2021).

115. E. S. J. Arnér, Selenoproteins—What unique properties can arise with selenocysteine in place of cysteine? Exp. Cell Res. 316, 1296–1303 (2010).

116. A. M. Diamond, et al., Dietary selenium affects methylation of the wobble nucleoside in the anticodon of selenocysteine tRNA([Ser]Sec). J. Biol. Chem. 268, 14215–14223 (1993).

117. U. Schweizer, S. Bohleber, W. Zhao, N. Fradejas-Villar, The Neurobiology of Selenium: Looking Back and to the Future. Front. Neurosci. 15, 652099 (2021).

118. X. Zhou, et al., Selenium Metabolism in Drosophila. J. Biol. Chem. 274, 18729–18734 (1999).

119. G. V. Kryukov, et al., Characterization of Mammalian Selenoproteomes. Science 300, 1439–1443 (2003).

120. J. Panee, Selenoprotein H Is a Redox-sensing High Mobility Group Family DNA-binding Protein That Up-regulates Genes Involved in Glutathione Synthesis and Phase II Detoxification*. Bind. Protein 282 (2007).

121. G. Li, et al., Gene expression profiling of selenophosphate synthetase 2 knockdown in Drosophila melanogaster. Metallomics 8, 354–365 (2016).

122. S. Blanco, et al., Aberrant methylation of t RNA s links cellular stress to neuro-developmental disorders. EMBO J. 33, 2020–2039 (2014).

123. J. Létoquart, et al., Insights into molecular plasticity in protein complexes from Trm9-Trm112 tRNA modifying enzyme crystal structure. Nucleic Acids Res. 43, 10989–11002 (2015).

124. S. J. Gratz, et al., Highly Specific and Efficient CRISPR/Cas9-Catalyzed Homology-Directed Repair in *Drosophila*. Genetics 196, 961–971 (2014).

125. K. Zhang, et al., An intellectual disability-associated missense variant in TRMT1 impairs tRNA modification and reconstitution of enzymatic activity. Hum. Mutat. 41, 600–607 (2020).

126. D. Su, et al., Quantitative analysis of ribonucleoside modifications in tRNA by HPLC-coupled mass spectrometry. Nat. Protoc. 9, 828–841 (2014).

